# Membrane Lipid Composition and Asymmetry Act as a Switch for Cell-Penetrating Peptide Translocation versus Leakage

**DOI:** 10.64898/2026.05.21.726760

**Authors:** Eric Catalina-Hernandez, Mateo Calle-Velasquez, Mohsen Habibnia, Marcel Aguilella-Arzo, Ramon Barnadas-Rodríguez, Victor A. Lorenz-Fonfria, Alex Peralvarez-Marin

## Abstract

Cell-penetrating peptides (CPPs) can enable intracellular access while avoiding cytotoxicity, yet their behavior is highly sensitive to membrane composition. Here we show that lipid composition and leaflet asymmetry act as a switch that determines whether the amphipathic CPP MAP undergoes non-disruptive translocation or stabilizes membrane pores that drive leakage. Using computational electrophysiology (CompEL) simulations in membranes of increasing physiological relevance, we find that symmetric anionic bilayers favor MAP insertion coupled to transmembrane pore stabilization in POPC:POPG membranes, reproducing progressive dye release in liposome leakage assays. Incorporation of cholesterol reduces the number of inserted peptides yet enhances pore stabilization, consistent with faster leakage kinetics in cholesterol-containing vesicles. In contrast, an asymmetric membrane model mimicking the eukaryotic plasma membrane (POPC outer leaflet; POPC:POPS inner leaflet) supports MAP translocation without sustained membrane disruption. This delivery-relevant mechanism is supported by efficient MAP internalization in HEK293 cells while maintaining high viability. Polarized ATR-FTIR provides structural context, indicating predominantly membrane-associated α-helical conformations across lipid compositions. Together, these results establish lipid asymmetry and composition as actionable biointerface parameters that tune CPP function between translocation and leakage and demonstrate an experimentally benchmarked framework for predicting membrane outcomes across complex lipid environments.

## 1. INTRODUCTION

Cell-penetrating peptides (CPPs) are short, cationic peptides capable of interacting with and translocating across membranes without causing cytotoxicity ^[1–3]^ and can be coupled to cargoes for intracellular delivery ^[4,5]^. Understanding the physicochemical basis of their translocation mechanisms (including lipid perturbation, pore formation, and direct penetration) is key to improving the design of CPPs and membrane-active peptides more broadly ^[6]^. However, translocation events occur on nanoscopic length scales and very short timescales, making transient intermediate and mechanistic details experimentally difficult to resolve. Molecular dynamics (MD) simulations complement experiments by providing atomistic insight into conformational transitions, peptide-lipid interactions, and membrane perturbation, while allowing systematic variation of parameters such as lipid composition, transmembrane potential, or peptide orientation.

Since CPP translocation occurs over seconds to minutes ^[7]^, conventional MD (cMD) cannot access these timescales directly, and enhanced sampling techniques are required ^[8]^. Methods such as steered MD ^[9,10]^, umbrella sampling ^[11,12]^, weighted ensemble ^[13,14]^, and adaptive steered MD combined with cMD^[15]^ can resolve translocation energetics and membrane disruption, but are computationally demanding, biased, or not readily accessible as entry-level approaches. Computational electrophysiology (CompEL) offers a practical alternative: by imposing an ion imbalance (ΔQ) that generates a transmembrane potential, it promotes membrane disruption in the form of transient pores or water channels that peptides can exploit to insert or translocate, without predefined reaction coordinates or forced sampling ^[16]^.

In our previous CompEL study ^[16]^, we screened five CPPs of varying physicochemical character, Arg9 (cationic), MAP and TP10 (amphipathic), and TP2 and K-FGF (hydrophobic), in POPC membranes across different ΔQ values and peptide numbers. We established that ΔQ 16 with eight peptides represents the optimal conditions for discriminating peptide-mediated membrane disruption. However, that study was limited to symmetric POPC membranes, preventing assessment of how lipid composition and bilayer asymmetry modulate CPP behavior.

Here, we extend the CompEL framework to membranes of increasing physiological complexity, POPC:POPG, POPC:POPG:CHOL, and asymmetric POPC:POPS, using MAP as a model CPP (Table 1), selected for its intermediate physicochemical properties and direct comparability with our previous work. Computational results are benchmarked against liposome leakage assays, FTIR structural characterization, and HEK293 cell internalization assays. This study aims to establish CompEL as a robust, accessible computational framework for resolving how membrane composition governs CPP mechanism, and for quantitatively correlating simulation outcomes with experimental observables.

**Table 1.**
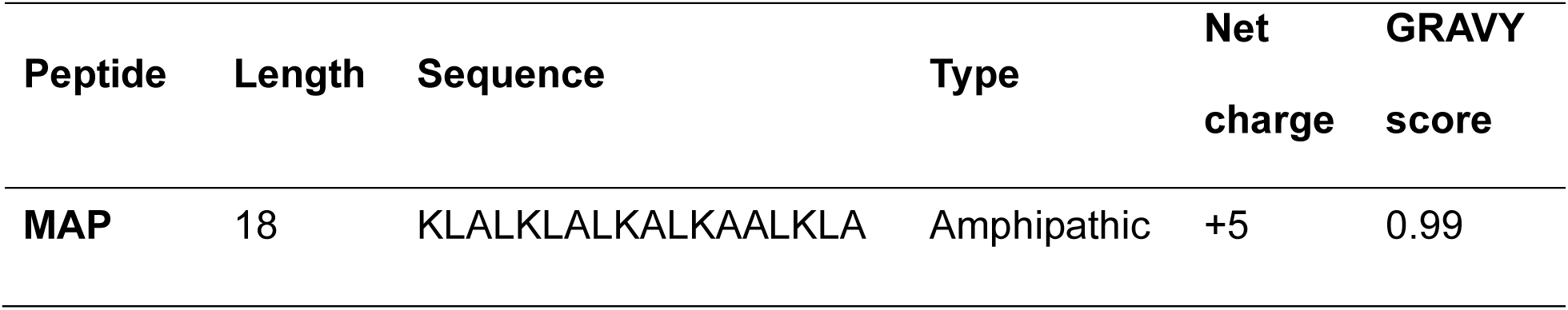
Sequence and characteristics of the peptide used in this study. GRAVY score is calculated using the Kyte-Doolittle scale^[33]^.

## 2. COMPUTATIONAL METHODS

### 2.1. Peptide simulation

As a model CPP, we used the MAP peptide ^[18,19]^ (Table 1) as in our previous study^[16]^. Briefly, MAP was modelled using AlphaFold 3^[20]^. Then, the peptide was introduced in a box of 7.5×7.5×7.5 nm and solvated with TIP3P waters. The system was minimized during 5 000 steps, equilibrated in the NVT ensemble during 125 000 steps, and a 250 ns production was run. CHARMM36m force field was selected. GROMACS^[21–28]^ software was used, specifically employing GROMACS 2020.7 package. The temperature was kept at 350 K throughout the study with the aim of accelerating the system dynamics. Periodic boundary conditions (PBC) were applied.

Clustering analysis of the peptide 250 ns trajectory was performed using MDAnalysis package in Python^[29,30]^. Later, the obtained centroid structure was used as input for the CompEL simulations.

### 2.2. Membrane systems set-up

Three membrane compositions were used. First, a symmetric membrane composed of POPC (1-Palmitoyl-2-oleoyl-sn-glycero-3-phosphatidylcholine), and POPG (1-Palmitoyl-2-oleoyl-sn-glycero-3-phosphatidylglycerol), using a 7:3 POPC:POPG mol ratio, namely POPC:POPG. Second, a symmetric cholesterol-containing membrane using a 6:3:1 POPC:POPG:CHOL mol ratio, namely POPC:POPG:CHOL. Third, an asymmetric membrane with POPC and POPS (1-Palmitoyl-2-oleoyl-sn-glycero-3-phosphatidylserine), with only POPC in the extracellular leaflet, and POPC:POPS with a 7:3 mol ratio in the intracellular leaflet, namely POPC:POPS membrane. Specific lipid compositions for all three membranes are displayed in Table 2. Systems without ions were built using CHARMM-GUI^[31–33]^ web server, solvating with TIP3P water. Ions were added in the CompEL set-up part.

**Table 2.**
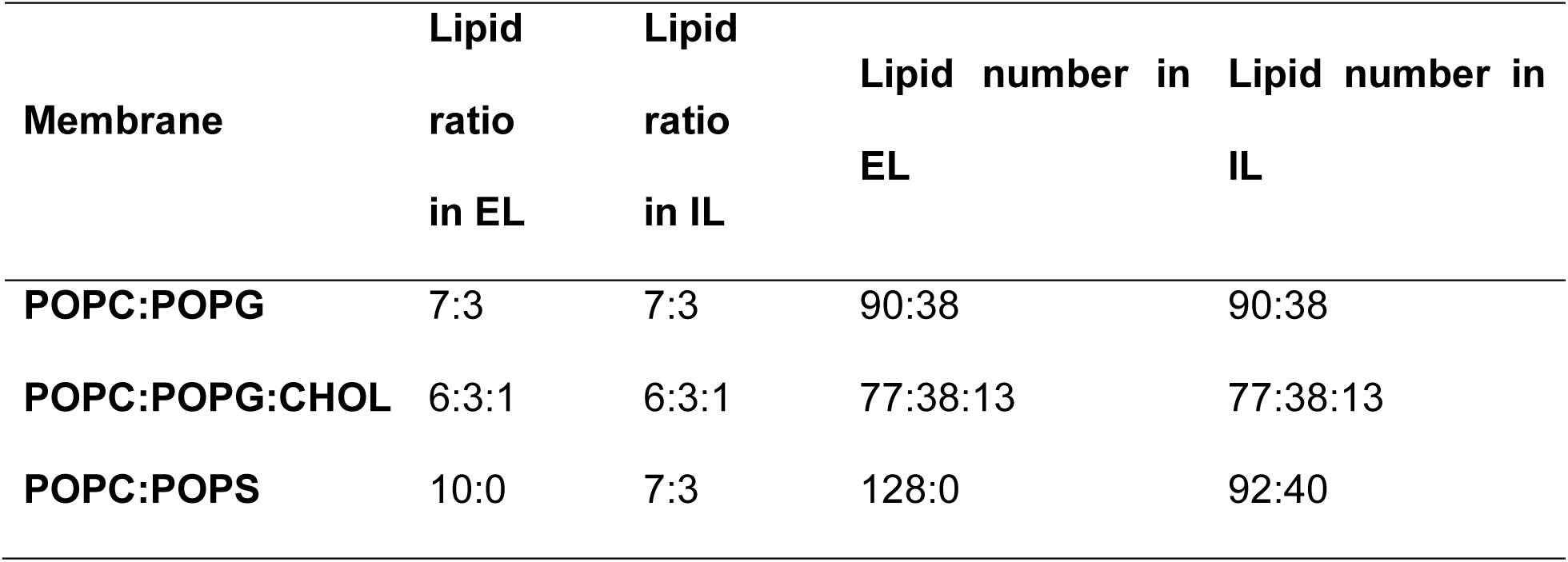
Membrane compositions used in the MD simulations of this study. Extracellular and intracellular leaflets are referred to as EL and IL, respectively.

### 2.3. CompEL set-up and simulations

CompEL involves the generation of transmembrane potential through ion imbalance, ΔQ, between both sides of the membrane. However, since PBC are applied, a double membrane configuration is required. To generate the desired transmembrane potential between one side and the other of the membrane, membrane systems without ions, obtained with CHARMM-GUI server, were used as input. The system was then duplicated, the second system was rotated, the box size was doubled, and both system files were concatenated into a single box with a double membrane configuration, as described in our previous study. Then, *gmx insert-molecules* utility was used to insert K^+^/Cl^−^ ions. The total number of ions includes those required to neutralize the charge of POPG/POPS lipids, and those required to generate ΔQ 16 (see Table 3). As can be seen, the goal is to obtain a total net charge of +8 in extracellular space, and −8 in the intracellular space (see Figure 1 for clarification). Simulations are run without peptides (control simulations) or with 8 MAP peptides. In the latter case, peptides are inserted in the extracellular space, along with the necessary counterions (5 per MAP peptide).

**Figure 1.**
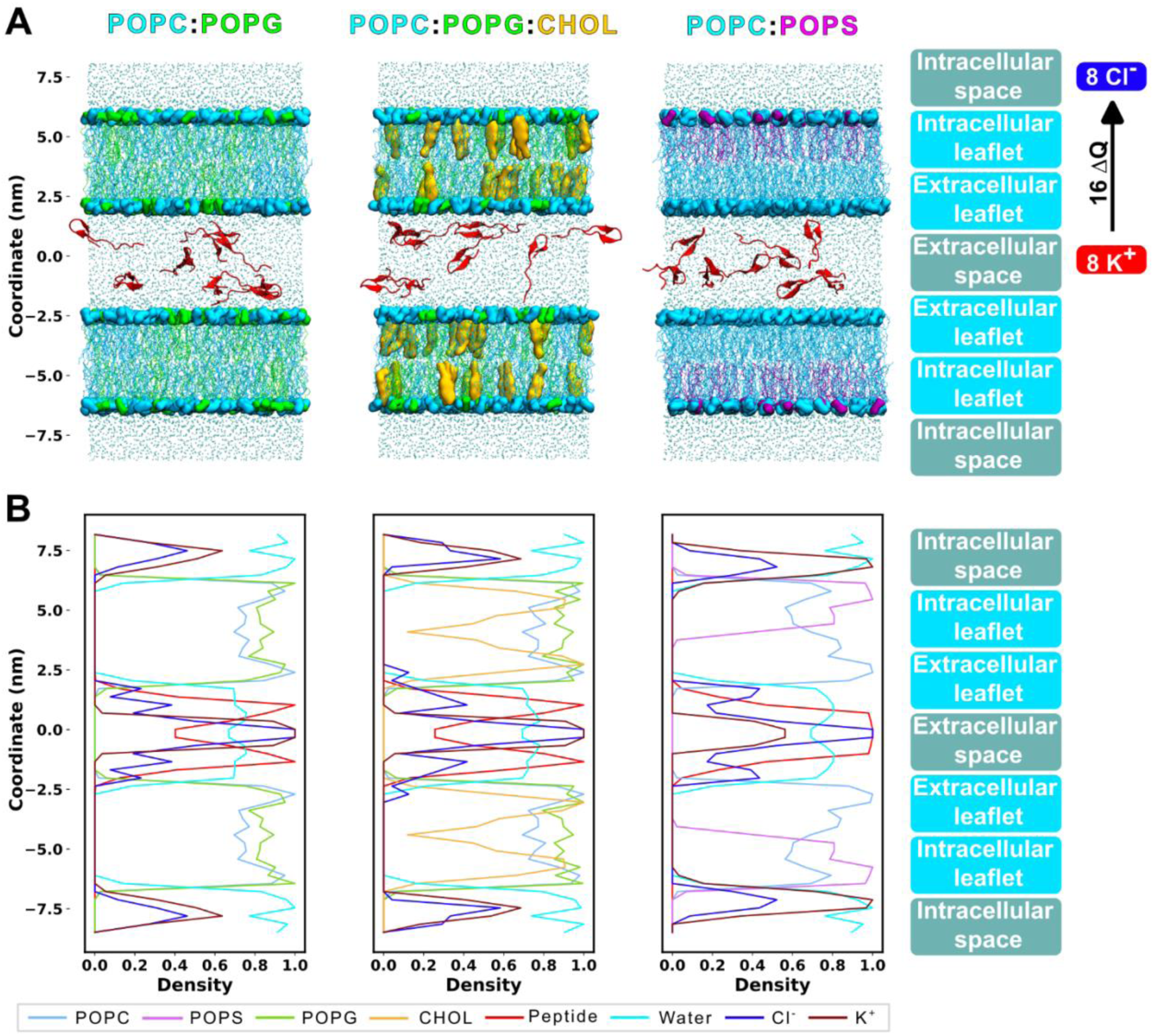
System set-up. **A.** Molecular representation of the starting point of the membranes. The polar heads for POPC (colored in light blue), POPG (in green), and POPS (in purple) are shown as QuickSurf, whereas lipid tails are represented as lines. Cholesterol (yellow) is shown as QuickSurf, and the peptides (red) are represented as NewCartoon. Water residues are shown as licorice (cyan). Extracellular and intracellular leaflets for both membranes are indicated. Peptides starting point in all simulations is the extracellular water compartment. The transmembrane gradient by charge imbalance (ΔQ) is represented indicated the 8 positive charges (through addition of K^+^ ions) in the extracellular water compartment and the 8 negative charges (addition of Cl^−^ ions) in the intracellular water compartment. **B.** Electron density analysis. The density values have been normalized for each individual species.

**Table 3.**
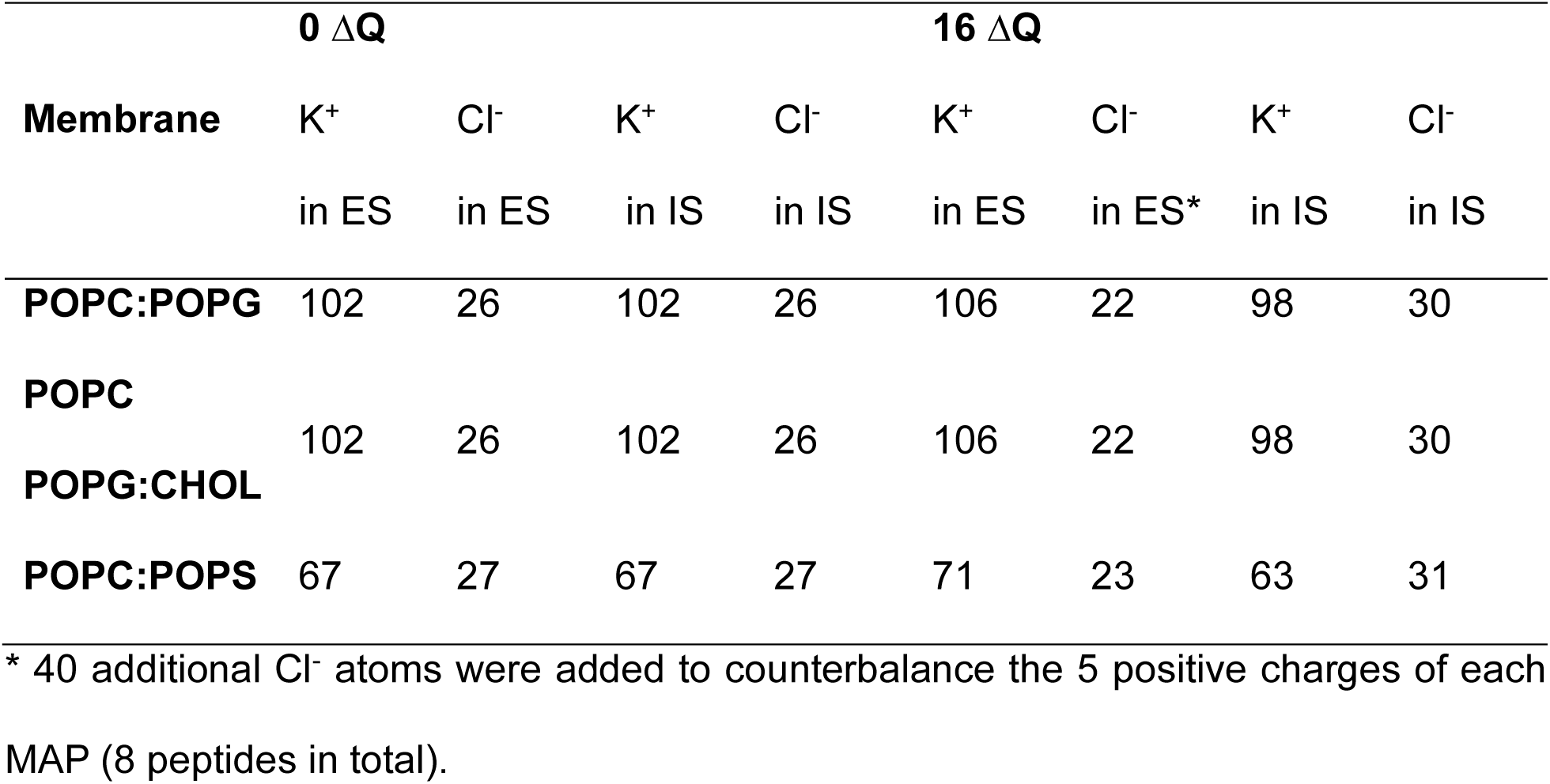
Total number of ions added in the CompEL simulations. Set-up with 0 ΔQ is shown to observe the difference between 0 and 16 ΔQ configuration. A 16 ΔQ refers to a net charge of +8 in the extracellular space, and a net charge of −8 in the intracellular space. In ΔQ 0, no net charge between spaces is present. Intracellular and extracellular spaces are referred to as IS and ES, respectively.

CompEL systems were minimized during 5 000 steps and equilibrated for ca. 2 ns. The equilibration procedure was run in six steps of 125, 125, 125, 500, 500, 500 ps, while lowering the positional restraints in each step: 1 000, 400, 400, 200, 40, 0 kJ·mol^−1^·nm^−2^, respectively. Finally, equilibrated systems were simulated during 500 ns, with 3 replicas for each system. Computational electrophysiology protocol^[34,35]^ was used in order to control ion/water position exchanges (all production files have been uploaded to the GitHub repository). Peptides and lipids were modeled using the CHARMM36m force field. Long-range electrostatics were calculated using the particle mesh Ewald (PME) method, using a real-space cutoff of 1.2 nm, and van der Waals interactions were force-switched between 1.0 and 1.2 nm using a Verlet cutoff scheme. All simulations have been conducted at 350 K. A total of 4.5 µs have been run for MAP simulations. Three replicas of control membranes (without peptides) have also been run, totaling 3 µs for control simulations, and 7.5 µs in total. Simulations have been run in a workstation with a GPU RTX3080Ti, at approximately 70 ns per day.

### 2.4. CompEL simulation analysis

Simulation analysis was performed in a Jupyter Notebook integrated development environment (IDE)^[36]^. Matplotlib^[37]^ was used for figure plotting. MDAnalysis^[29,30]^ was used to analyze system compositions, RMSD, and membrane thickness. Thickness was calculated for each simulation frame as the distance between the mean positions of the phosphate groups in the two membrane leaflets. *gmx* utilities were used to calculate electron density, potentials, and H bonds. PyLipID^[38]^ was employed to calculate occupancy, which indicates the percentage of simulation time that a peptide residue is in contact with a lipid. VMD^[39]^ was used for visual plotting and to analyze secondary structure using STRIDE^[40]^. The angle of the secondary structure of the peptide has been analyzed using the method presented by Choe^[41]^: i) calculate the center of masses of the first three and the last three residues involved in the α-helix, ii) find the vector connecting these two centers of masses, and iii) compute the angle between this vector and a unit vector parallel to the normal of the membrane. Thus, this script calculates the angle of the peptide with regard to the membrane normal when it adopts a helical structure. Consequently, a value close to 90° indicates that the peptide helix is perpendicular to the membrane normal, i.e., parallel to the membrane plane. Conversely, values closer to 0° indicate a peptide helix aligned with the membrane normal, i.e., perpendicular to the membrane plane. In the absence of helicoidal secondary structure, the script has been manually fine-tuned to calculate the representative orientation of the peptide. An in-house Python script based on Scipy^[42]^ was developed to calculate the pore radius. The membrane was divided along the bilayer normal (z-axis) into consecutive stacks 2 Å thick. For each stack and each simulation frame, the script identifies water molecules present within the membrane region and computes the maximum lateral distance between them. This distance is then used as a measure of the local pore, from which the corresponding pore radius is obtained. The analysis was performed over all simulation frames. The order parameter of C-H bonds of lipids (S_CH_), a parameter that characterizes the lipid order, was calculated using Equation 1:

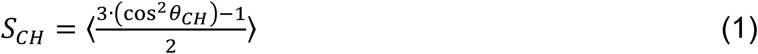

In this context, θ denotes the angle between the C-H bond of an acyl chain and a reference axis (typically the z axis in MD simulations). Angular brackets indicate an ensemble average over time and over all lipid molecules. The analysis was restricted to atoms belonging to the palmitoyl chain segment. Briefly, C and H atoms in the palmitoyl segment of POPC lipids were selected, the cosine of θ was computed and substituted into Equation 1, and the obtained values averaged between lipids and frames. In lipid membranes the value of the S_CH_ is negative, and so it is often reported as -S_CH_ in NMR and MD studies^[43]^. Average values are calculated by averaging over all C atoms. All analysis scripts are available in our GitHub repository (see section 2.9). Video S1 was produced using DaVinci Resolve software.

### 2.5. Liposome leakage experiments

POPC (Affymetrix, California, USA), POPG (Avanti, Alabama, USA), and cholesterol (Sigma-Aldrich, Missouri, USA) were dissolved in a chloroform/methanol mixture (2:1, v/v) to prepare lipid solutions with the following compositions: POPC:POPG (7:3 molar ratio), and POPC:POPG:CHOL (6:3:1 molar ratio). Liposomes were prepared as previously described ^[26]^. Briefly, the organic lipid mixtures were evaporated under reduced pressure to form a thin film. This film was hydrated with 10 mM phosphate buffer (pH 7.2) containing 2 mM of the fluorescent dye 8-hydroxypyrene-1,3,6-trisulfonic acid (HPTS), yielding multilamellar vesicles at a total lipid concentration of 10 mM. The vesicles were subsequently downsized by sequential extrusion through polycarbonate membranes with pore sizes of 800, 400, 200, and 100 nm. Dynamic light scattering (DLS, Nanotrac Wave, Microtrac, USA) revealed average radii of 90 nm for both POPC:POPG and POPC:POPG:CHOL liposomes. Non-encapsulated HPTS was removed by size-exclusion chromatography (SEC) using Sephadex G-25 PD-10 columns (Amersham Biosciences). Purified liposome samples were then supplemented with the quencher p-xylene-bis-pyridinium bromide (DPX, Fisher Scientific) to final concentrations of 5 µM DPX and 100 µM total lipid. The HPTS-loaded, DPX-containing liposomes containing DPX were transferred into black, clear-bottom 96-well plates. Fluorescence was recorded over time using excitation and emission wavelengths of 420 nm and 520 nm, respectively, on a FLUOstar Optima microplate reader (BMG LABTECH, Germany). The effect of MAP (Table 1 for sequence, purchased from SynPeptide, China) was assessed by adding them to the wells at a final concentration of 5 mM. After 4,000 s, complete fluorescence quenching was achieved by solubilizing liposomes with Triton X-100 (Sigma-Aldrich) at a final concentration of 5 µM. Data shown represents the average of three independent experiments.

### 2.6. Internalization and viability assays

HEK293 human cell lines were seeded in 24-well plates at a density of 200,000 cells per well and incubated for 48 hours prior to treatment. Cells were then exposed to triplicate TAMRA (5(6)-Carboxytetramethylrhodamine; Novabiochem®, Merck/Sigma-Aldrich, Cat. No. 851030) and TAMRA-labeled MAP at a final concentration of 1 µM for 1.5 hours. Following treatment, cells were harvested by trypsinization, washed and resuspended in flow cytometry buffer (PBS supplemented with 5% Fetal Bovine Serum). Cells were stained with 1ug/ml DAPI (ref: D9542(Merck)).

Flow cytometric analysis was performed using a CytoFLEX LX flow cytometer (Beckman Coulter), acquiring 10,000 events per sample. Doublets and aggregates were excluded by gating on forward scatter height versus area (FSC-H vs FSC-A) and side scatter width versus height (SSC-W vs SSC-H), respectively. Cell viability was assessed using DAPI staining, detected with excitation at 405 nm and emission collected at 450/45 nm. TAMRA fluorescence was detected using excitation at 561 nm and emission collected at 610 nm. Data were analyzed using CytExpert software (Beckman Coulter), and TAMRA signal was quantified as median fluorescence intensity (MFI) in viable, single-cell populations.

### 2.7. FTIR

The different lipids (POPC, POPG, POPS and CHOL, Avanti, Alabama, USA) were dissolved in chloroform/methanol (2:1, v/v) and mixed to obtain the following compositions: 100% POPC, 7:3 POPC:POPG, 7:3 POPC:POPS, 6:3:1 POPC:POPG:CHOL (molar ratios). The different lipid solutions were dried in a rotavapor at 158 mbar for 30 min and then resuspended in the peptide solution (250 μM TAMRA-labeled MAP in 10 mM phosphate buffer at pH 7.2) to a final MAP concentration of 50 μM and a mol lipid/peptide ratio of 20. Then they were vigorously vortexed to obtain multilamellar vesicles (MLVs).

Samples for polarized ATR-FTIR experiments were obtained by adding 4 μL of the different MLVs suspension containing the peptides on top of a 4 mm diameter diamond ATR crystal (3 active reflections DuraDisk, Czitek) and letting them slowly dry at ambient humidity to form oriented multilayer membranes. Once dried, they were hydrated by adding on top of them a drop (6 μL) of a 200 mM NaCl solution at pH 7.2 (high ionic strength was used to keep the hydrated multilayer membranes close to the ATR crystal surface).

FTIR spectra were obtained with a Vertex80 FT-IR spectrometer (Bruker). Interferograms of sample and sample-free (reference) ATR crystals were acquired with a photovoltaic MCT detector and a mobile mirror speed corresponding to a 40 kHz modulation of the built in HeNe laser, later processed to obtain absorbance spectra at 2 cm^−1^ spectral resolution extending from 4000 to 800 cm^−1^. Polarized spectra were acquired using IR light polarized parallel (Abs_0_) and perpendicular (Abs_90_) to the plane of incidence, produced with a BaF_2_ holographic wire grid polarizer (Thorlabs, WP25H-B) placed before the sample and mounted on a motorized rotational stage (Thorlabs, PRM1/MZ8). Interferogram acquisition was synchronized with the rotation of the polarizer using TTL pulses, obtaining polarized spectra by repeatedly alternating 0° and 90° polarization and averaging the results. To analyze the polarized spectra of hydrated membranes we first subtracted the dominant liquid water contribution (O-H stretching band around 3300 cm^−1^), using polarized spectra of the buffer, recorded in the same ATR system.

Dichroic ratios (R^ATR^) of bands of interests, mainly for the lipid CH_2_ symmetric band and the peptide amide I band, were calculated by dividing their integrated area between Abs_0_ and Abs_90_ spectra. The dichroic ratio of a band can be related to the second order parameter of the angle of the transition dipole moment of the vibration responsible for the band with respect to the ATR surface normal (S_α_), and to the three-dimensional components of the relative intensity of the evanescent electric field at the ATR crystal-sample interface^[27]^:

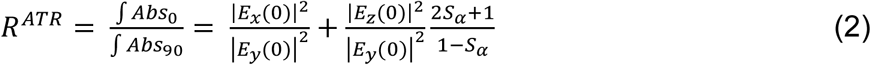

Then, the value of S_α_ for vibration can be obtained from the experimentally determined value of R^ATR^ for a band as the values of |*E_x_*(0)|^2^, |*E_y_*(0)|^2^ and |*E_z_*(0)|^2^. These last values were calculated using the Harrick thick-film weak-absorber approximation^[27]^, the crystal and sample refraction index (2.39 and 1.40, respectively) and the angle incidence if the IR beam (60°). For the lipids, the analysis of the symmetric CH_2_ vibration of the methyl acyl groups gives us the averaged order parameter of the bisection of the two C-H methyl bonds, which can be compared with the averaged S_CH_ obtained by MD simulations. For the amide I the value of averaged S_α_ turned to be less useful, as the amide I band turned to be heterogenous, displaying different structural components (see Results).

Structural information about MAP was obtained by decomposing the amide I band into a sum of Voigtian bands (convolution of Lorentzian and Gaussian bands). The process involves first band-narrowing and Fourier self-deconvolution (FSD), allowing to better resolve underlaying components, followed by band decomposition using curve-fitting^[28–30]^, both steps done using homemade scripts running in MATLAB using a narrowing factor of 2 and a Lorentzian width of 18 cm^−1^. For accurate results, this decomposition was conducted on isotropic spectra (Abs_iso_), free from orientation effects, obtained from the experimentally measured polarized spectra (Abs_0_ and Abs_90_) as^[31]^:

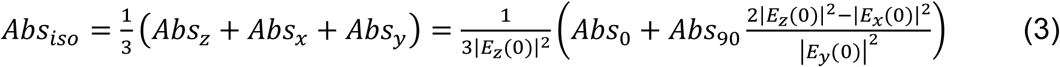

The area of the resulting fitted bands was assigned to different secondary structures based primary on their wavenumber value^[32]^, but also on their width and on their intensity and sign in the corresponding linear dichroism spectrum (see below).

### 2.8. Linear dichroism

Given the complexity of the amide I band of the peptide, with many subcomponents, we used the linear dichroism (LD) spectra rather than the dichroic ratio to obtain orientation information. The LD spectrum was obtained as:

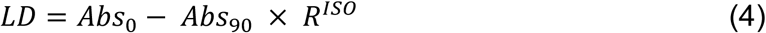

where Abs_0_ and Abs_90_ correspond to the spectrum obtained at 0° and 90° polarization, respectively. R^ISO^ is the so-called isotropic dichroic ratio, the value of R^ATR^ expected for a band with S_α_ = 0 (non-preferent orientation), which in our setup was 1.78. In a LD spectrum the intensity (and area) of a band becomes proportional to the value of S_α_ of the vibration responsible for it: bands from vibrations with S_α_ > 0 (parallel to the surface normal) will appear as positive, those from vibrations with S_α_ < 0 (perpendicular to the surface normal) will appear as negative bands, and those from vibrations with S_α_ = 0 will be cancelled out. Because for helices the amide I vibration has its TDM roughly along the helical axis, a transmembrane helix will give a positive band in the LD spectrum, while a helix oriented in the plane of the membrane is expected to give a negative band in the LD spectrum. Amide I bands from peptides in solution or adopting a random coil structure are expected to be missing in the LD spectrum. We applied FSD to LD spectra to enhance the resolution of bands in the amide I region.

### 2.9. Data availability

Simulation inputs, and analysis code can be found at our GitHub repository: https://github.com/APMlab-memb/CompEL_MAP.git

Due to file size limitations, the simulation trajectory file will be shared upon request.

## 3. RESULTS

One conclusion of our previous computational study on CPP-mediated membrane disruption using CompEL simulations^[16]^ was that the most effective discriminating conditions are a transmembrane potential of ΔQ 16 with eight peptides. However, that study was limited to symmetric POPC membranes, which contrasts with the complex, asymmetric lipid composition of biological membranes where CPPs operate. Here, we extend the CompEL framework to bilayers of increasing physiological complexity (incorporating negatively charged lipids, cholesterol, and membrane asymmetry) and directly compare simulation outcomes with experimental data.

Computationally, three replicas of 500 ns CompEL ΔQ 16 simulations were performed in symmetric POPC:POPG (7:3) and POPC:POPG:CHOL (6:3:1) membranes, and in an asymmetric POPC:POPS membrane (POPS restricted to the intracellular leaflet at a 7:3 POPC:POPS ratio) as a simplified eukaryotic plasma membrane model (Figure 1). The simulation system consists of two lipid bilayers separated by two aqueous compartments representing the extracellular (positive membrane potential) and intracellular spaces (Figure 1). Eight MAP peptides (P:L 1:32) were placed in the extracellular compartment at the start of each simulation.

For clarity, the bilayer leaflet facing the extracellular compartment is referred to as the extracellular leaflet, and the leaflet facing the intracellular compartment as the intracellular leaflet (Figure 1). Three peptide behaviors are distinguished: (i) partitioning: transition from the extracellular aqueous phase to the water-bilayer interface; (ii) insertion: deeper penetration into the hydrophobic core, with contacts established with both acyl tails and intracellular leaflet headgroups; and (iii) translocation: complete bilayer crossing to reach the intracellular aqueous compartment. In all conditions, only one or two peptides insert or translocate, while the remainder partition into the extracellular leaflet, consistent with previous reports ^[51]^. Analysis therefore focuses on peptides exhibiting insertion or translocation.

Consistent with our previous work ^[16]^, the applied transmembrane potential induces transient pore formation in peptide-free control simulations. These pores are short-lived and close rapidly, yielding small average pore sizes (Table S1), characteristic of electroporation ^[52,53]^. Pore formation is thus primarily driven by the ionic imbalance, and these transient defects serve as permissive intermediates that MAP peptides can exploit to initiate stabilized, peptide-mediated membrane disruption. The following sections examine how MAP interacts with and modulates these transient pores across membrane compositions of increasing physiological complexity.

### 3.1. POPC:POPG

CompEL ΔQ 16 simulations in POPC:POPG (7:3) membranes show that the most frequent outcome is the insertion of two MAP peptides, while the remaining six partition into the extracellular leaflet (Figure 2A, Video S1, Table S2). Peptide insertion stabilizes the transient pores induced by the ionic imbalance, expanding their average radius relative to peptide-free controls (Tables 4 and S1). Conversely, the replica in which all peptides remain partitioned displays the smallest average pore size (Table 4), confirming that insertion, not partitioning, drives pore stabilization and expansion.

**Figure 2.**
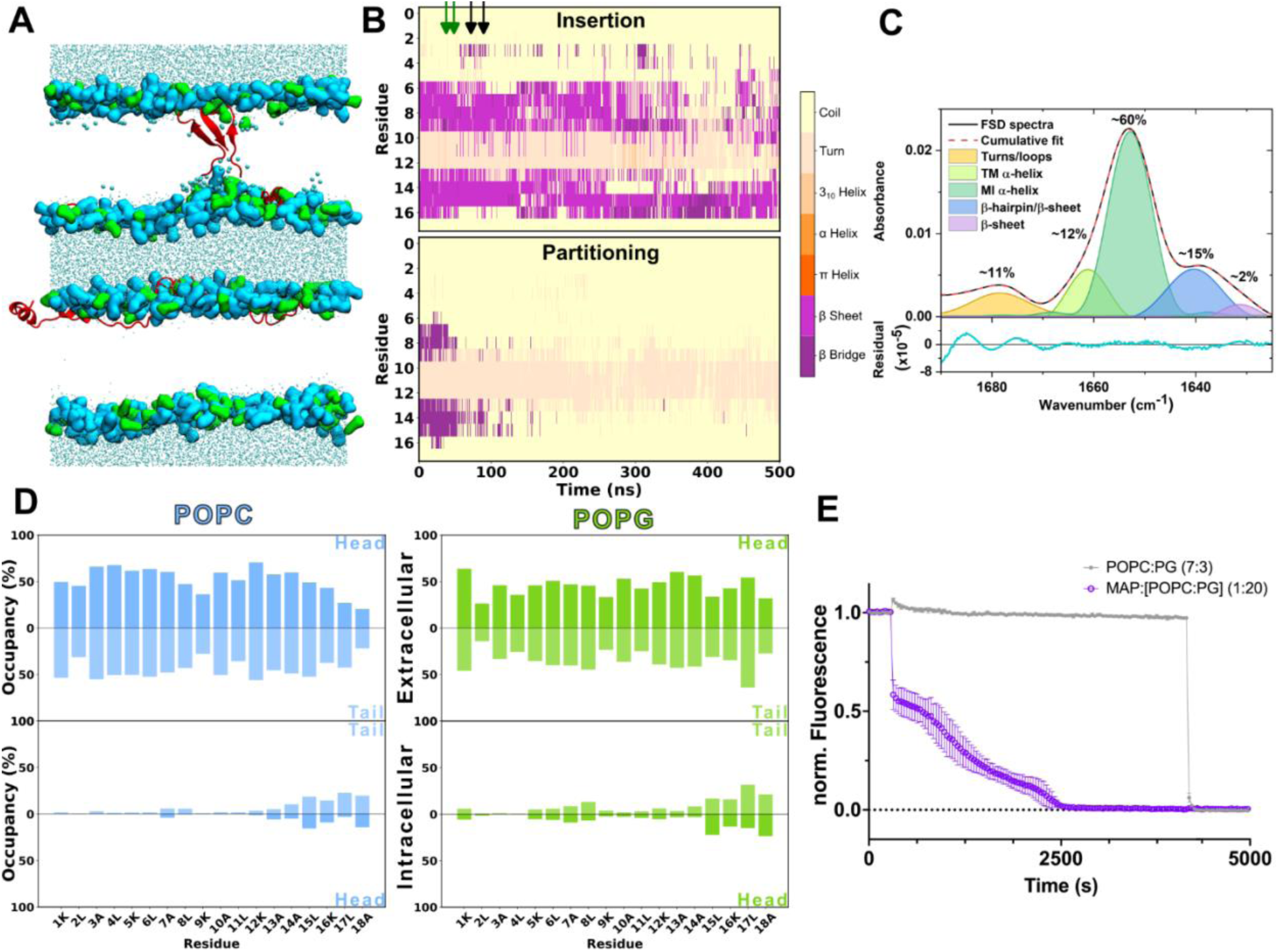
POPC:POPG CompEL 16 ΔQ 500 ns simulation with 8 MAP peptides results. **A.** Representative molecular configuration at the end of the 500 ns CompEL simulation. In two of the three replicas, two peptides achieve insertion, whereas the other six peptides are partitioning with the inner leaflets. Lipid polar heads are represented in QuickSurf and colored in light blue (POPC) or green (POPG). Peptides are represented as NewCartoon and colored in red. Water molecules are shown as licorice and colored in cyan, whereas larger water residues are used for the water molecules in the pore. Lipid tails are omitted for clarity. The molecular dynamics simulation process is presented in Video S1. **B.** Secondary structure evolution throughout the simulation. The average between the three replicas is shown, differentiating between peptides that insert or partition. The arrows indicate the moment where insertion occurs: the green arrows refer to the two insertions in the second replica, whereas the black arrows represent the insertion of the two peptides in third replica. **C.** Average secondary structure estimation by amide I band decomposition of band-narrowed FTIR spectra. The peptide adopts mainly a membrane interface α-helical structure (∼60 %, green band), with few contributions of turns/loops (∼11 %, yellow band), transmembrane α-helix (∼12 %, light green band), and β-hairpins (∼15 %, blue band), with just around 2 % adopting a β-sheet conformation (purple band). The fitting residual was in the order of 10^−5^. The MI helix can be mixed with random coil contributions. **D.** Average residue occupancy by POPC (left) or POPG (right). The occupancy is differentiated between extra and intracellular leaflets, and between lipid heads (darker blue) or tails (lighter blue). **E.** Liposome leakage assays monitored by HPTS fluorescence quenching. Fluorescence traces are shown for liposomes in the absence of peptide (control, grey) and after addition of MAP (5 µM, purple).

**Table 4.** Average pore radius (Å) for the 3 membranes, differentiating by replica, and indicating the standard deviation.

| Replica | POPC:POPG | POPC:POPG:CHOL | POPC:POPS |
| --- | --- | --- | --- |
| 1 | 0.28 ± 0.02 | 2.02 ± 0.20 | 2.21 ± 0.23 |
| 2 | 3.10 ± 0.23 | 5.05 ± 0.48 | 1.80 ± 0.20 |
| 3 | 3.25 ± 0.27 | 3.80 ± 0.31 | 1.95 ± 0.21 |

In computational secondary structure analysis, all peptides begin with β-sheet conformation. Upon membrane interaction, most lose this structure; however, inserted peptides retain β-sheet conformation throughout, likely reflecting timescale limitations (Figure 2B). Membrane interaction is further supported by an increase in peptide-lipid H-bonds and a concomitant decrease in peptide-water H-bonds, indicating the transition of MAP from bulk solution to the water-bilayer interface (Figure S1A).

FTIR spectroscopy was used to characterize the secondary structure of MAP in POPC:POPG membranes. The amide I band reflects a time-averaged signal across the full MAP population, and band narrowing reveals multiple components indicative of marked structural heterogeneity (Figure 2C). Two bands appear in the α-helical region: a band at 1654 cm⁻¹ (60% of total amide I area), assigned to a membrane-interface (MI) helix based on its wavenumber and negative LD signal (Figure S2), and a band at 1660 cm⁻¹ (12%), assigned to a transmembrane (TM) helix based on its high wavenumber, consistent with a highly hydrophobic environment, and positive LD signal (Figure S2). A band at 1643 cm⁻¹ (15%), absent in the LD spectrum, lacks a clear structural assignment: its wavenumber is too low for a soluble helix yet too high for a canonical parallel β-sheet ^[54]^, and is tentatively attributed to a parallel β-sheet or β-hairpin, the latter expected near 1640–1645 cm⁻¹ in H₂O based on model peptide data in D₂O ^[55]^. A minor band at ∼1634 cm⁻¹ (2%) is consistent with β-sheet conformation; if arising from an antiparallel β-sheet, as observed in MD simulations, a companion band at 1685–1695 cm⁻¹ (∼1/10th the intensity) would be expected but is likely masked by the more intense band at ∼1680 cm⁻¹ (11%), which is assigned to turns and loops, though coiled-coil α-helical structures represent an alternative assignment ^[56]^. Taken together, the amide I profile indicates that MAP in POPC:POPG membranes is predominantly membrane-associated, mainly as MI helices, with a smaller TM helix population and minor β-sheet and turn contributions.

Residue-level lipid occupancy analysis (Figure 2D) shows that most peptides interact with both POPC and POPG headgroups and acyl chains in the extracellular leaflet, while inserted peptides extend their contacts to the intracellular leaflet. Despite a 7:3 POPC:POPG molar ratio, occupancy levels for both lipids are comparable, indicating a preferential interaction of MAP with the anionic POPG lipids.

Liposome leakage experiments (Figure 2E) confirm that MAP at 5 µM induces progressive fluorescence loss in POPC:POPG (7:3) liposomes, consistent with membrane disruption and dye release driven by pore formation.

### 3.2. POPC:POPG:CHOL

In POPC:POPG:CHOL (6:3:1) simulations, one MAP peptide inserts per replica while the remaining seven partition into the extracellular leaflet (Figure 3A, Video S1, Table S2). As in POPC:POPG membranes, transient pores form in peptide-free controls but close rapidly (average pore size 0.3 Å, Table S3). Peptide insertion stabilizes these pores, yielding average pore sizes of 2–5 Å (Table 4), comparable to those in POPC:POPG membranes. Notably, the cholesterol-containing membrane supports the insertion of only one peptide per replica versus two in POPC:POPG, suggesting that cholesterol moderately reduces MAP’s membrane-disruptive capacity.

**Figure 3.**
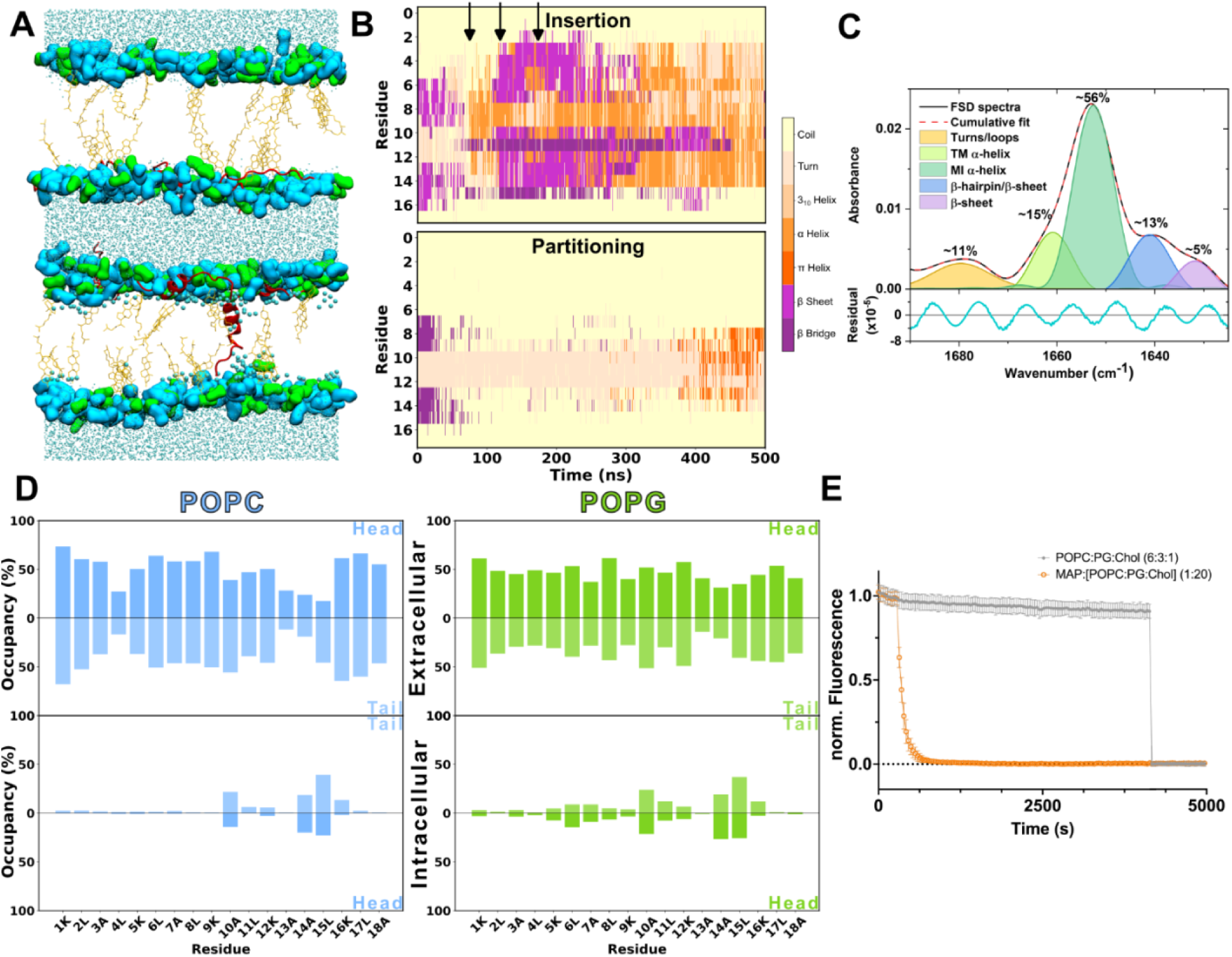
POPC:POPG:CHOL MAP CompEL ΔQ 16 500 ns simulation results. **A.** Representative molecular configuration of the CompEL system at the end of the simulation. In all three replicas, one peptide is inserted, whereas the remaining seven peptides are partitioning. The peptides are shown as cartoon and colored in red, the polar heads are shown as QuickSurf and colored in light blue (POPC) or green (POPG), cholesterol lipids are shown as orange licorice, and water residues as cyan licorice. Larger water residues are used for the water molecules in the pore. Lipid tails are omitted for clarity. The molecular dynamics simulation process is presented in Video S1. **B.** Secondary structure of the 8 MAP peptides during the 500 ns of CompEL simulation, differentiating between insertion and partitioning behaviors. The arrows indicate the moment where insertion occurs. Each arrow represents the insertion in one of the replicas. **C.** Average secondary structure estimation from FTIR experiments. MAP adopts mainly a MI α-helical structure (∼56 %, green band), with few contributions of: turns/loops (∼12 %, yellow band), TM α-helix (∼15 %, light green band) and β-hairpins (∼14 %, blue band), with just around 5 % adopting a β-sheet conformation (purple band). The fitting residual was in the order of 10^−5^. The MI helix can be mixed with random coil contributions. **D.** Occupancy of the peptide residues by POPC or POPG lipids. The occupancy is differentiated between extra and intracellular leaflets, and lipid head or tails. Cholesterol occupancy is shown in Figure S1C. **E.** Results of liposome leakage experiments. Fluorescence levels compare liposomes without peptide (control, grey), and with peptide addition (5 µM, orange).

In this membrane, β-sheet structure is lost upon partitioning, whereas inserted peptides rapidly acquire α-helical conformation (Figure 3B), a more pronounced structural conversion than observed in POPC:POPG. Peptide-lipid H-bond formation and reciprocal loss of peptide-water contacts confirm membrane association (Figure S1B).

FTIR analysis of MAP in POPC:POPG:CHOL membranes (Figure 3C) reveals a band distribution closely resembling that of POPC:POPG membranes. The MI α-helix is again the dominant conformation (1652 cm⁻¹, 56%, negative LD signal), followed by TM helix (1661 cm⁻¹, 15%, positive LD signal). Unoriented components include a band at ∼1641 cm⁻¹ (13%, tentatively assigned to parallel β-sheet or β-hairpin), a band at 1631 cm⁻¹ (5%, antiparallel β-sheet), and a band at 1680 cm⁻¹ (11%, turns/loops). The overall FTIR profile is consistent with that of POPC:POPG membranes, suggesting that the addition of 10 mol% cholesterol does not substantially alter the bulk secondary structure distribution of MAP.

Peptide-lipid occupancy (Figure 3D) mirrors the pattern observed in POPC:POPG membranes, with comparable POPC and POPG interaction levels and no prominent cholesterol contacts (Figure S1C), further highlighting the primacy of electrostatic interactions with anionic lipids in MAP membrane association.

Liposome leakage experiments (Figure 3E) show that MAP induces rapid and pronounced fluorescence loss in POPC:POPG:CHOL vesicles, indicative of fast membrane disruption and dye release, consistent with pore-mediated leakage.

### 3.3. POPC:POPS

To better approximate the asymmetric lipid composition of the eukaryotic plasma membrane, a bilayer model with POPS restricted to the intracellular leaflet was employed. POPS was selected over POPG for its higher physiological relevance^[57]^. Since asymmetric liposomes cannot be readily formed, MAP internalization was assessed in human cells alongside CompEL asymmetric membrane simulations. FTIR experiments were performed using symmetric POPC:POPS membranes which, despite the inherent limitation of membrane symmetry, serve as a representative model of the intracellular leaflet composition to characterize MAP interactions with negatively charged phospholipids.

In asymmetric POPC:POPS simulations, MAP translocation, not only insertion, is observed in two independent replicas (Table S2), a behavior distinct from the POPC:POPG and POPC:POPG:CHOL conditions. As in previous compositions, a transient pore is generated by the ion imbalance (Table S1), which MAP exploits to insert and stabilize, but pore sizes (∼2 Å, Table 4) are smaller than in POPC:POPG and POPC:POPG:CHOL, suggesting that translocation is coupled to pore closure rather than sustained pore expansion (Table 4). The selective enrichment of anionic POPS lipids in the intracellular leaflet electrostatically drives MAP toward that leaflet, facilitating full bilayer crossing (Figure 4A, Video S1).

**Figure 4.**
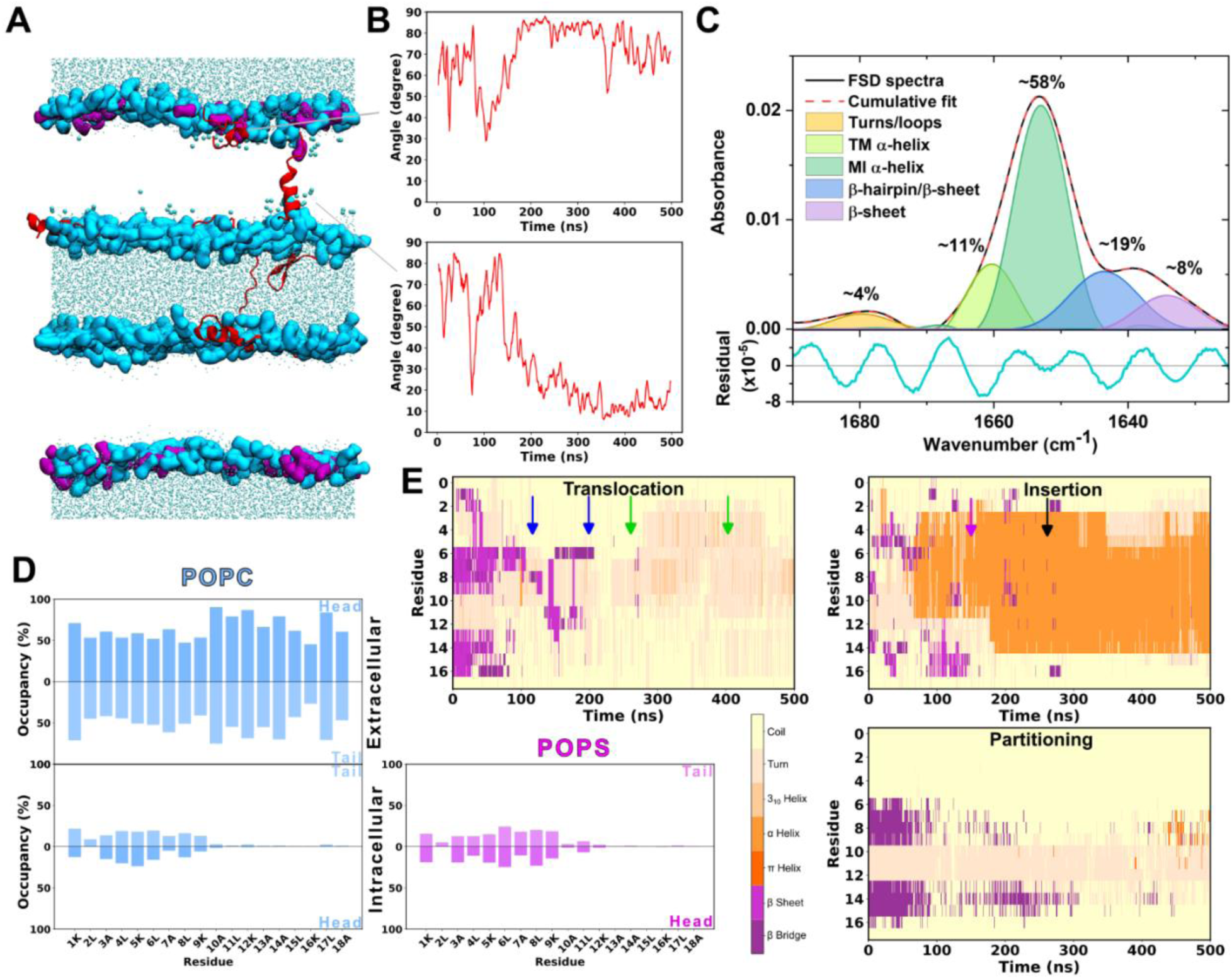
POPC:POPS MAP CompEL ΔQ 16 500 ns simulation results. **A.** Representative molecular configuration of the POPC:POPS CompEL system, with one insertion and one translocation. Polar heads are shown as QuickSurf in light blue (POPC) or purple (POPS). Peptide is shown as NewCartoon in red. Water molecules are shown as licorice in cyan, with larger water residues for the water molecules present in the pore. The molecular dynamics simulation process is presented in Video S1. **B.** Evolution of the peptide angle throughout the simulation. The peptide that translocates (upper plot) and the peptide that gets inserted (lower) are shown. **C.** Average secondary structure estimation from FTIR experiments. The majority of the peptide adopts a MI α-helical structure (∼58 %, green band), with few contributions of turns/loops (∼4 %, yellow band), TM α-helix (∼11 %, light green band) and β-hairpins (∼19 %, blue band), with just around an 8 % adopting a β-sheet conformation (purple band). The fitting residual was in the order of 10^−5^. The MI helix can be mixed with random coil contributions. **D.** Average residue occupancy by POPC and POPS lipids. **E.** Secondary structure throughout the simulation. The average among the three replicas is shown, differentiating by peptide behavior. The arrows indicate the moment where insertion/translocation occurs. In the translocation plot, each color represents one peptide, the first arrow indicating the moment of insertion, and the second one the translocation. In the insertion plot, each color represents the moment of insertion of one of the peptides.

Throughout translocation, MAP undergoes a well-defined reorientation (Figure 4B), quantified using the approach of Choe ^[41]^. Upon partitioning into the inner leaflet, the peptide lies parallel to the bilayer plane (∼90°). During insertion, it reorients perpendicular to the membrane axis (∼0°), consistent with prior reports ^[51]^. Upon completing translocation into the outer leaflet, it returns to a parallel orientation (∼90°), mirroring the partitioning geometry.

FTIR analysis (Figure 4C) confirms that the MI helix remains the dominant conformation in POPC:POPS membranes (1653 cm⁻¹, ∼58%, negative LD signal), with a TM helix accounting for ∼11% of the amide I area (1660–1661 cm⁻¹, positive LD signal). Unoriented components include a band at 1644 cm⁻¹ (∼19%, soluble α-helix or parallel/hairpin β-sheet), a band at 1634 cm⁻¹ (∼8%, antiparallel β-sheet), and a band at ∼1680 cm⁻¹ (∼4%, turns or coiled-coil), all showing weak or absent LD signals. The FTIR profile is broadly consistent across all three membrane compositions tested, indicating that the bulk secondary structure of MAP is relatively insensitive to the specific anionic lipid identity (POPG vs. POPS), while the simulation data reveal critical differences in translocation outcome driven by membrane asymmetry.

Peptide-lipid occupancy analysis (Figure 4D) shows that MAP interaction in the extracellular leaflet is restricted to POPC, the sole lipid present there. In the intracellular leaflet, POPS and POPC display comparable occupancy values despite POPS being the minority lipid species, underscoring the importance of electrostatic interactions in driving MAP toward anionic lipids.

Computational secondary structure analysis (Figure 4E) reveals that MAP initially adopts β-sheet conformations in solution, reflected by higher peptide-peptide versus peptide-lipid H-bonds (Figure S1D). Upon membrane contact, β-sheet structure dissolves: partitioning peptides adopt predominantly turn and coil conformations, while inserting peptides acquire α-helical structure. Translocated peptides remain largely in coil and turn conformations throughout the simulation, with only a minor and transient helical component emerging at later timepoints, suggesting that translocation occurs without requiring a stable helical intermediate.

Internalization and viability assays in HEK293 cells (Figure S4) confirm that MAP efficiently enters human cells, with uptake approaching 100% and cell viability remaining above 90%, consistent with CPP behavior and supporting the translocation mechanism identified in the asymmetric POPC:POPS simulations.

### 3.4. Membrane Biophysical Parameters

Electric field and electrostatic potential profiles for POPC:POPG (Figure S5A), POPC:POPG:CHOL (Figure S6A), and POPC:POPS (Figure S7A) membranes are consistent with those reported in previous computational studies [53,58,59]. Membrane thickness and lipid order parameters show no significant differences across compositions (Figures S5B, S6B, S7B), indicating that the structural integrity of the bilayer is preserved across conditions.

FTIR-derived lipid order parameters, reflecting average CH₂ orientation across both acyl chains, and MD-derived order parameters are compared in Table S3. Both approaches yield values in a comparable range, consistent with previous studies ^[43]^, and indicate that MAP-membrane interaction does not globally perturb bilayer order regardless of membrane composition or peptide behavior. This validates the structural integrity of the simulation systems and supports the reliability of cross-composition comparisons.

### 3.5. Composition-Dependent MAP Behavior: A Cross-System Comparison

Integrating computational and experimental data across all four membrane compositions, namely POPC^[16]^, POPC:POPG, POPC:POPG:CHOL, and asymmetric POPC:POPS, reveals a clear and composition-dependent pattern in MAP behavior (Figure 5A, Video S1, Table S4).

**Figure 5.**
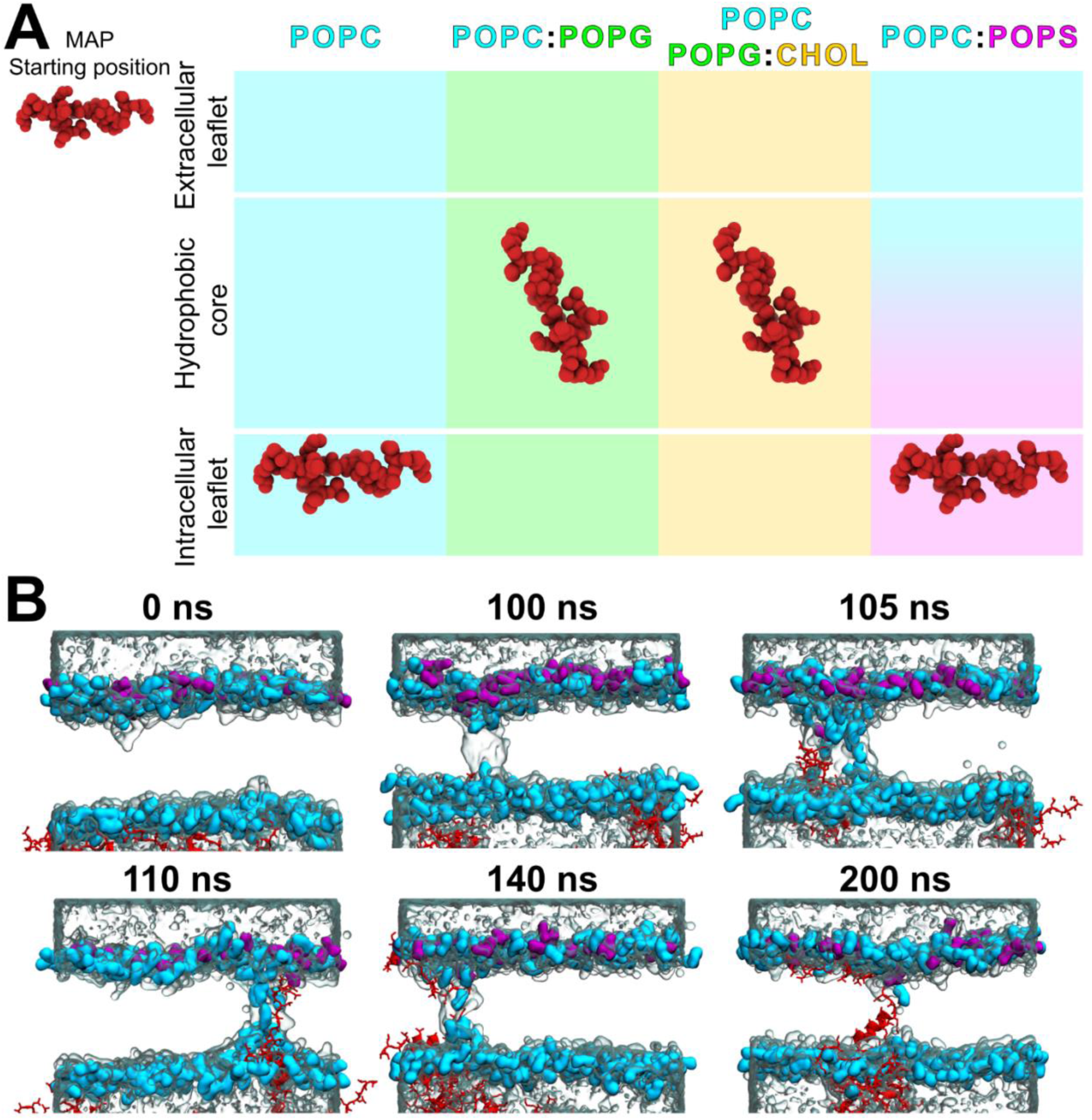
Peptide insertion and translocation. **A.** Comparison of MAP insertion extent across membranes. The results from four membranes are shown: POPC (from our previous study), POPC:POPG, POPC:POPG:CHOL, and POPC:POPS. The starting point is represented in the extracellular leaflet, with insertion in POPC:POPG, and POPC:POPG:CHOL, and translocation to the intracellular leaflet in POPC and POPC:POPS membranes. The molecular dynamics simulation process for all three membrane compositions is presented in Video S1. **B.** Process of pore formation, insertion, and translocation in POPC:POPS. The start of the simulation (0 ns), of the pore (100 ns), and of the insertion (105 ns), followed by full insertion (110 ns), start of the translocation and the insertion of another peptide (140 ns), and end of translocation (200 ns) steps are shown. Polar heads are represented as QuickSurf in light blue (POPC) or purple (POPG). The peptides are shown as NewCartoon, the side chain is shown as licorice and colored in red. Waters are shown as transparent QuickSurf and colored in cyan.

In POPC membranes, MAP translocates across the bilayer. The addition of anionic POPG to the extracellular leaflet abolishes translocation: MAP instead inserts and stabilizes membrane pores, with two peptides inserting per replica. Incorporation of cholesterol further reduces insertion to one peptide per replica, indicating that both extracellular anionic lipids and cholesterol progressively suppress MAP’s membrane-crossing capacity ^[60–64]^. Critically, when asymmetry is restored by confining POPS to the intracellular leaflet, as in biological membranes, translocation is recovered, driven by electrostatic attraction of MAP toward the negatively charged inner leaflet^[64]^. This composition-dependent mechanistic switch, from translocation to insertion and back to translocation, constitutes the central finding of this study.

Across all membrane compositions, FTIR experiments consistently show that MAP adopts predominantly a MI α-helical conformation (∼58–63%, 1652–1654 cm⁻¹), followed by unoriented components (soluble helix or β-sheet, ∼13–19%), TM helix (∼9–15%), and minor β-sheet (∼2–8%) and turn (∼4–12%) populations (Table S4, Figure S8A). POPC membranes alone produce a very slow liposome leakage (Figure S8B), consistent with rapid pore-closing translocation, whereas POPC:POPG and POPC:POPG:CHOL membranes show progressive and rapid leakage, respectively, reflecting sustained pore stabilization by inserted peptides.

The full mechanistic sequence of MAP translocation is best illustrated in the asymmetric POPC:POPS system, following the pore formation framework of Levine & Vernier ^[65]^ (Figure 5B). The applied ΔQ induces pore initiation (∼100 ns), followed by water infiltration and lipid headgroup rearrangement that drives pore maturation (∼105 ns). MAP inserts into the mature pore and adopts a perpendicular orientation (∼110 ns). Electrostatic interactions with POPS lipids in the intracellular leaflet then initiate translocation (∼140 ns), culminating in reorientation to a parallel configuration and full bilayer crossing (∼200 ns). Pore closure accompanies translocation, consistent with the lower average pore sizes observed in this system (Table 4). A second peptide inserts subsequently, driven by the amphipathic nature of MAP, paralleling our previous observations in POPC bilayers ^[16]^.

## 4. DISCUSSION

### 4.1. Membrane Composition Drives a CPP/AMP Mechanistic Switch in MAP

The central finding of this study is that MAP does not operate through a single, fixed mechanism. Instead, its mode of membrane interaction, translocation versus pore stabilization, is determined by lipid composition. In symmetric membranes containing anionic lipids in both leaflets (POPC:POPG, POPC:POPG:CHOL), MAP inserts and stabilizes transmembrane pores, behavior characteristic of antimicrobial peptides (AMPs) (Leontiadou et al., 2006). In membranes where anionic lipids are absent (POPC) or asymmetrically restricted to the intracellular leaflet (POPC:POPS), more closely approximating eukaryotic cell membranes, MAP translocates across the bilayer without sustained pore formation, consistent with CPP behavior. This composition-dependent functional plasticity, where the same peptide switches between AMP-like and CPP-like mechanisms, is an emerging principle in the field (Gehan et al., 2020; Pae et al., 2014; Sharmin et al., 2016) and is here directly demonstrated across four membrane systems with converging computational and experimental evidence.

The mechanistic basis of this switch lies in the interplay between electrostatics and membrane asymmetry. In symmetric anionic membranes, the extracellular leaflet presents a negatively charged surface that electrostatically traps MAP at the interface, promoting insertion and pore stabilization rather than crossing. Cholesterol further rigidifies the bilayer and reduces defect probability, explaining the lower insertion frequency observed in POPC:POPG:CHOL versus POPC:POPG ^[60–62]^. In contrast, in the asymmetric POPC:POPS membrane, the anionic gradient across the bilayer generates a directional driving force that pulls MAP toward the intracellular leaflet, favoring translocation and subsequent pore closure^[64,67,68]^. The recovery of translocation in the asymmetric system, and its loss in symmetric anionic membranes, directly implicates bilayer asymmetry, not merely anionic lipid content, as the key determinant of CPP behavior.

### 4.2. CompEL Simulations Capture a Pre-Pore Mechanism

The translocation and insertion events observed across all membrane compositions are consistent with the pre-pore model proposed for CPP membrane crossing^[69–70]^, which posits that transient nanoscale lipid packing defects, generated by membrane tension or electrostatic stress, serve as short-lived entry points for CPPs before resealing occurs^[71–73]^.

In our peptide-free control simulations, the applied CompEL ΔQ induces spontaneous, transient water defects that collapse rapidly, behavior consistent with pre-pore-like intermediates driven by electrostatic stress^[52,53]^. When MAP is present, these defects are captured and stabilized: peptides are attracted to the water channel, insert into the membrane, and either extend pore lifetime (AMP-like, in POPC:POPG systems) or exploit the defect transiently to translocate before pore closure (CPP-like, in POPC and POPC:POPS systems). The conductive pores observed in peptide-containing simulations thus represent electrostatically stabilized pre-pore analogues, whose fate, persistent or transient, is governed by membrane composition and the consequent peptide-lipid interactions.

It is important to acknowledge that while the applied transmembrane potentials fall within ranges used in established computational studies^[53,58,59]^, they substantially exceed physiological membrane potentials (0 to −150 mV) and experimental electroporation thresholds (∼0.5 V)^[74]^. These elevated potentials are a well-established strategy in atomistic MD to overcome timescale limitations and accelerate rare membrane remodeling events and should be interpreted as effective driving forces rather than direct representations of physiological conditions. Within this constraint, CompEL simulations provide mechanistic resolution of defect-mediated translocation and pore stabilization that is qualitatively consistent with experimentally proposed pre-pore mechanisms.

### 4.3. Convergence of Computational and Experimental Data

A strong quantitative and qualitative agreement is observed between CompEL simulations and experimental assays across all membrane compositions, supporting the ability of this approach to capture biologically relevant membrane-peptide interactions[34].

In POPC membranes, MAP induces a very slow liposome leakage (Figure S8B), consistent with rapid translocation and pore closure in the simulations, a relationship between translocation and membrane resealing supported by prior work^[68,71,72]^. In POPC:POPG membranes, MAP induces progressive leakage (Figure 2E), in agreement with simulation results showing peptide insertion and sustained pore stabilization with larger average pore sizes (Table 4). POPC:POPG:CHOL membranes show even faster leakage kinetics (Figure 3E), consistent with the formation of comparably sized or larger pores in the simulations (Table 4). Finally, in the asymmetric POPC:POPS system, simulated translocation is directly supported by near-complete MAP internalization in HEK293 cells with minimal cytotoxicity (Figure S4), confirming CPP behavior under physiologically relevant asymmetric conditions.

Peptide secondary structure shows consistent cross-validation between FTIR and computation. FTIR experiments across all compositions reveal that MAP is predominantly in a MI α-helical conformation (∼58–63%), with minor TM helix (∼9–15%), unoriented helix/β-sheet (∼13–19%), and turn (∼4–12%) populations (Table S4). In MD simulations, MAP initially adopts β-sheet conformations in solution; upon membrane interaction, these dissolve into turn/coil conformations for partitioning peptides or α-helical structure for inserting peptides. The partial and transient helicoidal structuring observed computationally at late simulation timepoints is consistent with the experimentally detected predominance of helical conformations on the experimental timescale (minutes vs. nanoseconds) and mirrors the membrane-driven helix stabilization reported for structurally related antimicrobial peptides such as Pa-MAP upon interaction with anionic vesicles^[75]^. Secondary structure evolution thus reflects a progressive, membrane-driven conformational transition from β-sheet in solution to helix at the membrane interface, captured at different points along this trajectory by MD and FTIR respectively.

Peptide orientation analysis further supports this picture. LD spectroscopy confirms that MAP adopts a predominantly interfacial (MI) orientation in all membrane compositions, while the computationally tracked angular reorientation during translocation (Figure 4B) provides mechanistic resolution of this average signal: partitioning (∼90°), insertion (∼0°), and post-translocation re-partitioning (∼90°) form a sequential reorientation trajectory^[41,51]^. Residue-level occupancy analysis reveals preferential interaction with anionic lipids (POPG, POPS) relative to POPC across all compositions, with C-terminal residues leading insertion in symmetric membranes and N-terminal residues gaining prominence in the asymmetric POPC:POPS system^[76–78]^, suggesting that the terminal residue driving membrane engagement is itself sensitive to lipid asymmetry.

Membrane lipid order parameters derived from FTIR and MD simulations are in comparable ranges across all compositions (Table S3), consistent with previous studies^[43]^, and confirm that MAP-membrane interaction does not globally perturb bilayer order regardless of peptide behavior. Differences in absolute values between methods are attributable to the fundamentally different timescales: MD probes nanosecond-scale local dynamics, while FTIR reflects ensemble-averaged properties over minutes, during which cumulative peptide-induced remodeling and relaxation can occur.

Finally, RMSD analysis and trajectory inspection indicate that systems reach overall structural stabilization within 250 ns (Figure S9), suggesting that this timescale is sufficient to discriminate peptide behaviors (insertion, translocation, orientation). Analysis of slower processes, such as secondary structure conversion, benefits from the full 500 ns trajectories used here, or ideally microsecond simulations where computationally feasible. The straightforward parallelizability of CompEL across replicas makes this approach computationally tractable compared to methods requiring single ultralong trajectories.

### 4.4. Mechanistic Implications and Limitations

The composition-dependent CPP↔AMP switch demonstrated for MAP likely reflects a general principle applicable to other membrane-active peptides whose mechanisms have been reported to vary with lipid environment^[61,62,65]^. From a therapeutic perspective, this plasticity implies that the in vivo efficacy and selectivity of MAP and related peptides will be strongly influenced by the specific lipid composition of target membranes, which varies between cell types, subcellular compartments, and pathological states. As shown in symmetric PS membranes, computational predictions indicate a preference for membrane partitioning over insertion (Figure S3A), a helical secondary structure preference for insertion (Figure S2 and S3B), and cholesterol-induced hindrance of membrane perturbation due to tighter packing (Figure S3C). These findings agree with the ability of MAP to translocate efficiently across an asymmetric membrane model resembling the eukaryotic plasma membrane while causing minimal membrane distortion and cytotoxicity (Figure S4), a characteristic particularly relevant for its potential as a selective intracellular delivery vehicle in eukaryotic cells.

A key methodological limitation of the CompEL framework is that the sustained transmembrane potential biases the system toward pore-mediated translocation pathways, rendering alternative mechanisms such as direct membrane permeation without pore formation, endocytic uptake, or large-scale lipid rearrangements, inaccessible within these simulations. The findings presented here should therefore be interpreted as evidence for a pore-mediated mechanism under high-field conditions, while not excluding the coexistence of additional translocation pathways under different physiological contexts. This constraint should be considered when extrapolating these mechanistic conclusions to in vivo systems.

## 5. CONCLUSIONS

This study expands the application of CompEL to membrane-active peptides, demonstrating its power to resolve composition-dependent mechanisms of peptide-membrane interaction and benchmarking simulation outcomes against complementary experimental data. The central finding is that MAP exhibits a composition-driven mechanistic switch: it behaves as a CPP, translocating across the bilayer in POPC and asymmetric POPC:POPS membranes, the latter serving as a simplified model of the eukaryotic plasma membrane, while adopting an AMP-like mode, inserting and stabilizing transmembrane pores in symmetric POPC:POPG and POPC:POPG:CHOL membranes. This switch is governed by the distribution of anionic lipids across leaflets: extracellular anionic lipids trap MAP at the interface promoting pore stabilization, whereas asymmetric restriction of anionic lipids to the intracellular leaflet generates a directional electrostatic driving force that favors translocation. Cholesterol further modulates this behavior by reducing insertion propensity. These computational findings are directly supported by liposome leakage assays and, critically, by MAP internalization in human cells with minimal cytotoxicity, validating the asymmetric membrane model as a physiologically relevant proxy.

Across all compositions, occupancy analyses confirm strong preferential interaction of MAP with anionic lipids (POPG, POPS), reflecting its cationic character, while secondary structure and orientation analyses consistently show progressive helix stabilization upon membrane association. The agreement between CompEL simulations and experimental observables, leakage kinetics, FTIR secondary structure, lipid order parameters, and cell internalization, establishes CompEL as a reliable, accessible, and computationally efficient framework for dissecting peptide-membrane interactions at mechanistic resolution.

Future work should extend this approach to membranes of greater biological complexity, incorporating phosphatidylethanolamine (PE), phosphatidylinositol (PI), more diverse asymmetric lipid distributions, and membrane protein components, to better recapitulate the cellular environment. We anticipate that CompEL will serve as a broadly applicable entry-level tool for investigating the membrane activity of peptides and other membrane-active molecules.

## AUTHOR CONTRIBUTIONS

Conceptualization: E.C.-H., and A.P.-M.; supervision: A.P.-M., and V.A.L.-F.; methodology and formal analysis: all authors; investigation: all authors; funding acquisition and resources: A.P.-M., V.A.L.-F., R.B.-R., M.A.-A; writing–original draft preparation: E.C.-H and A.P.-M.; writing–review and editing: all authors. All authors have read and agreed to the sent version of the manuscript.

## ACKNOWLEDGEMENTS

Authors acknowledge financial support by the Spanish Government Grant PID2020-120222GB-I00 (to A.P.-M.), PID2022-142795NB-I00 (to M.A.-A), PID2021-123682OB-I00 (to R.B.-R), PID2022-137582NB-I00 (to V.A.L.-F) funded by MCIN/AEI/10.13039/501100011033, and Universitat Autònoma de Barcelona predoctoral fellowship (B21P0033 to E.C.-H.), predoctoral fellowship PRE2020-093992 founded by MCIN/AEI/ 10.13039/501100011033 and by “ESF Investing in your future” (to M.C.-V.),and Universitat Jaume I project UJI-B2022-42 (to M.A-A), a Generalitat Valenciana project CIAICO/2023/106 (to A.P.-M. and M.A.-A.), and an Excellence Unit “María de Maeztu” grant CEX2019-000919-M founded by MCIN/AEI/ 10.13039/501100011033.

