## Supplementary Material for "Membrane Lipid Composition and Asymmetry Act as a Switch for Cell-Penetrating Peptide Translocation versus Leakage"

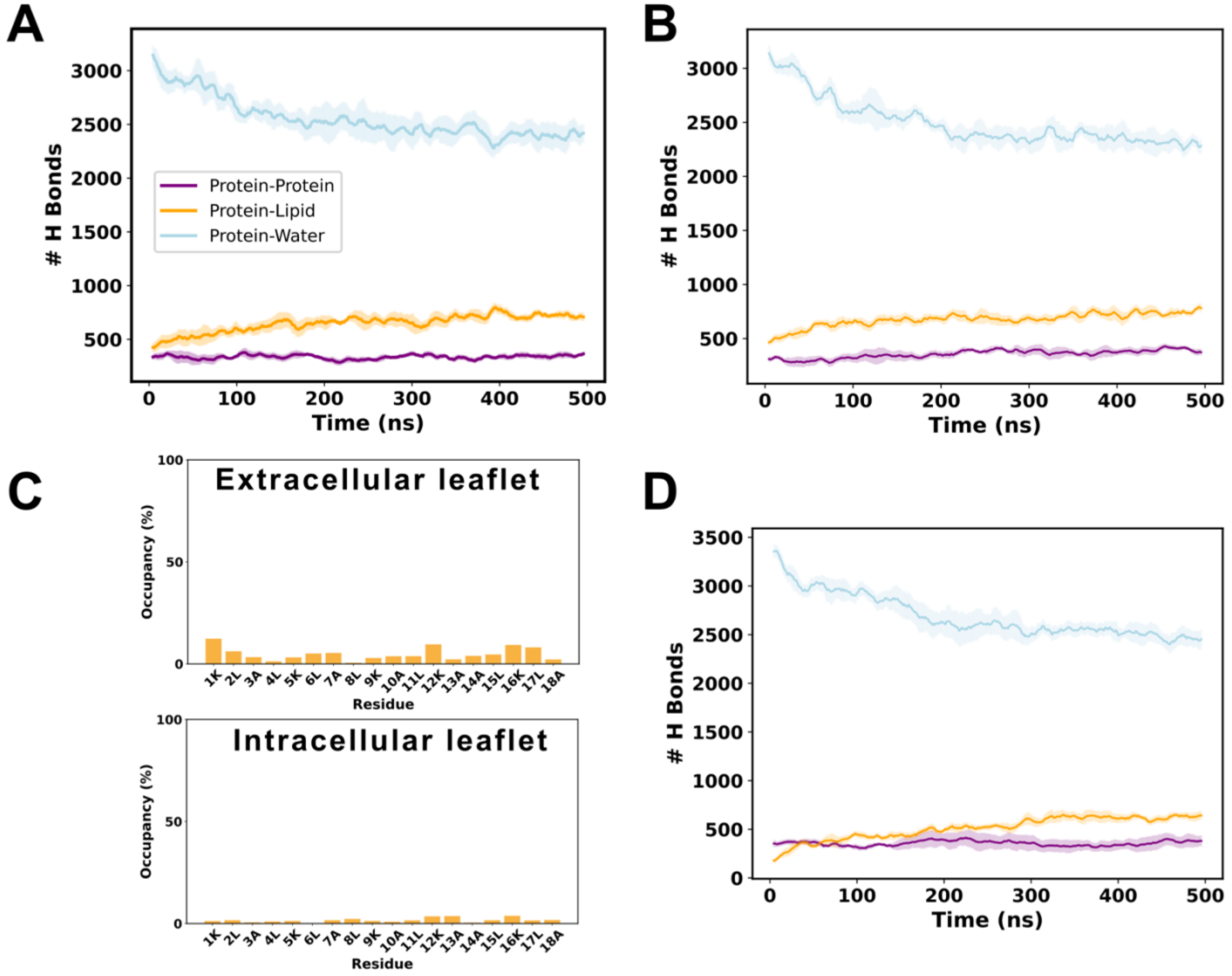


**Figure S1. A,B,D.** Average number of hydrogen bonds formed by peptides, lipids and waters throughout the simulation in POPC:POPG (A), in POPC:POPG:CHOL (B) and in POPC:POPS (D). **C.** Peptide residues occupancy by cholesterol in POPC:POPG:CHOL simulations. Occupancies are differentiated between upper and lower leaflets.

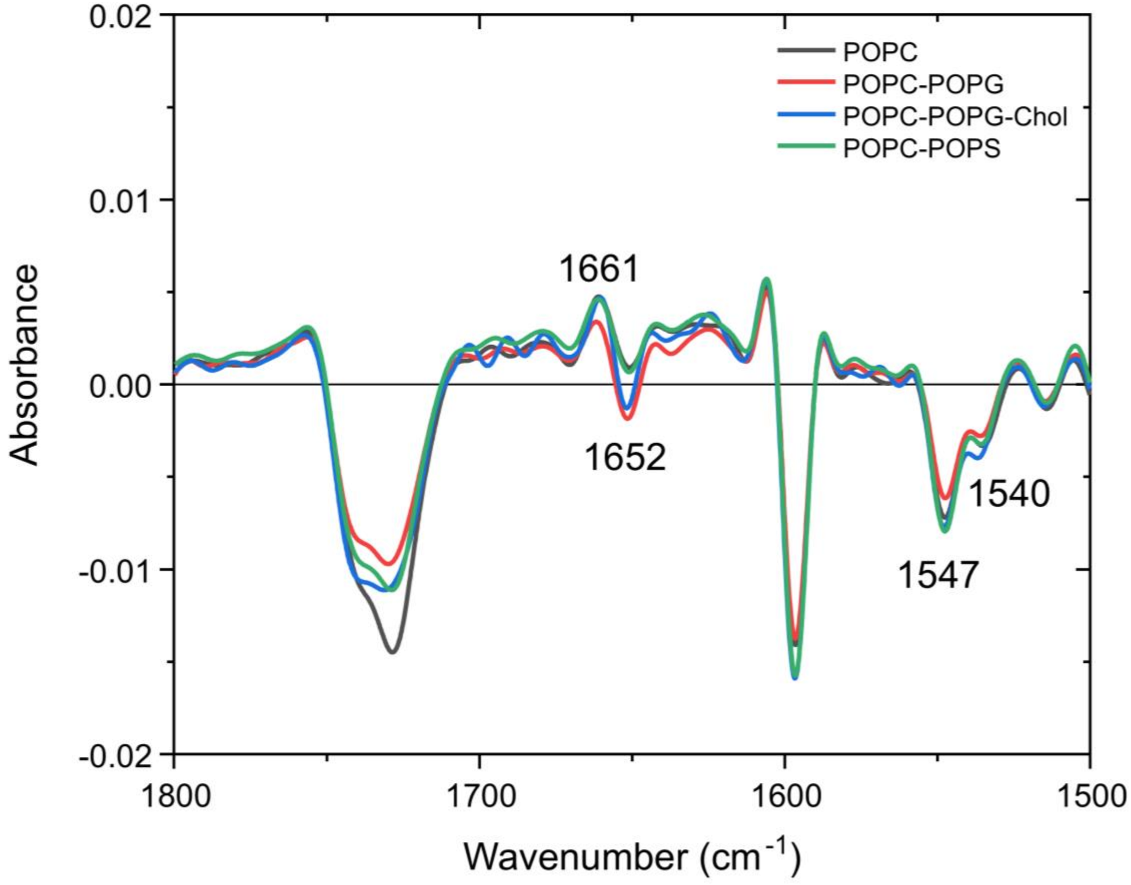


**Figure S2.** Linear dichroism spectra of the peptide MAP in the four membrane compositions: POPC (black), POPC:POPG (red), POPC:POPG:CHOL (blue), POPC:POPS (green). The LD was band-narrowed using FSD. In all cases, MAP exhibits a similar LD spectrum: a positive signal at 1661 $cm^{-1}$ paired with a negative signal at 1547 $cm^{-1}$, characteristic of a TM peptide. A second population presents a negative band and a downshift in the amide I region (1652 $cm^{-1}$) together with a positive, downshifted amide II band (1540 $cm^{-1}$), indicative of a MI conformation. It can also be seen ta negative signal around 1730 -1740 $cm^{-1}$ corresponding to C=O stretching from phospholipid's esters, and another signal around 1600 $cm^{-1}$ which can be assigned to the fluorescent TAMRA group attached to the peptide.

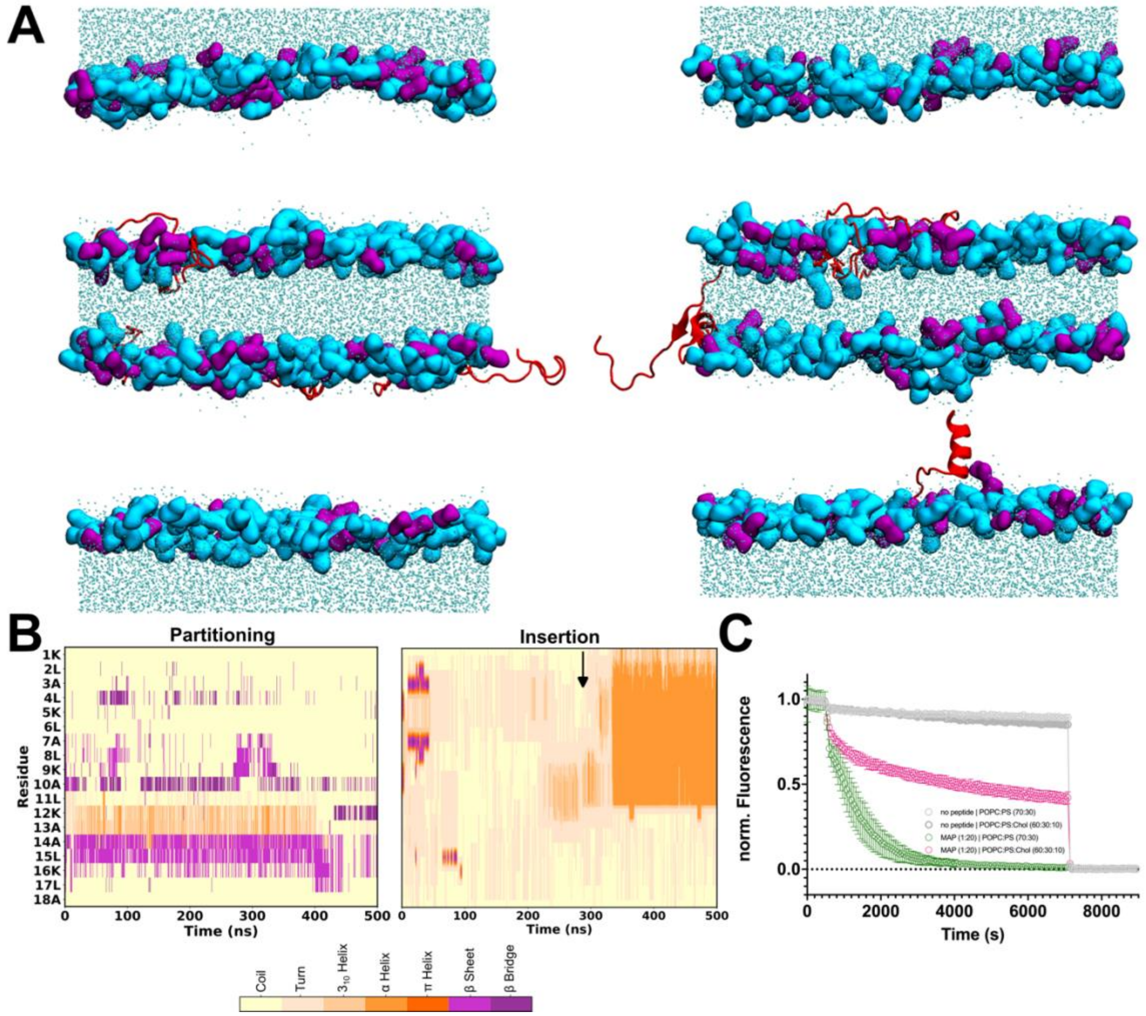


**Figure S3. A.** Molecular representation of the last frame of the simulation showing partitioning of all eight peptides (left, in two out of three replicas), and insertion into the membrane of one peptide and seven partitioning peptides (right, in one out of three replicas) in the system in the ΔQ 16 CompEL symmetric POPC:POPS simulation. The peptides are shown as cartoon and colored in red, the polar heads are shown as QuickSurf and colored in light blue (POPC) or green (POPG), cholesterol lipids are shown as orange licorice, and water residues as cyan licorice. Lipid tails are omitted for clarity. **B.** Secondary structure of MAP peptides during the 500 ns symmetric POPC:POPS simulation showing peptides that partitioned (left) and the peptide that got inserted into the membrane (right). **C.** Leakage experiments in symmetric POPC:POPS and POPC:PS:Chol liposomes in the absence and presence of MAP (n=3, mean±S.E.M).

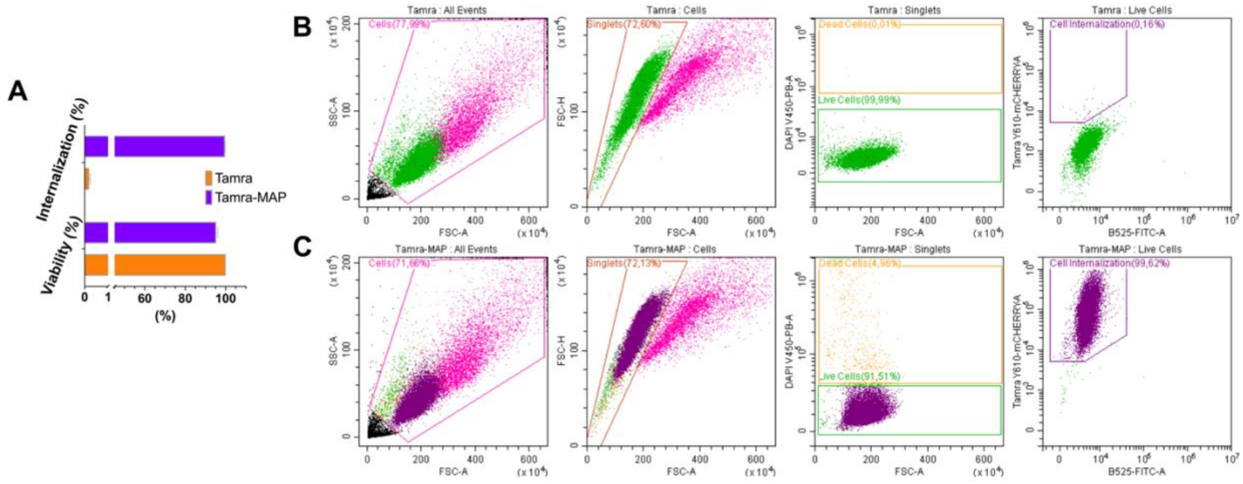


**Figure S4. A.** Results of HEK293 cells internalization and viability assays. Data compare untreated controls with samples exposed to MAP (n=3; mean±S.E.M.). **B.** Representative flow cytometry results for internalization and viability experiments of the Carboxytetramethylrhodamine (TAMRA) fluorescent dye. **C.** Representative flow cytometry results for internalization and viability experiments of MAP coupled with TAMRA dye.

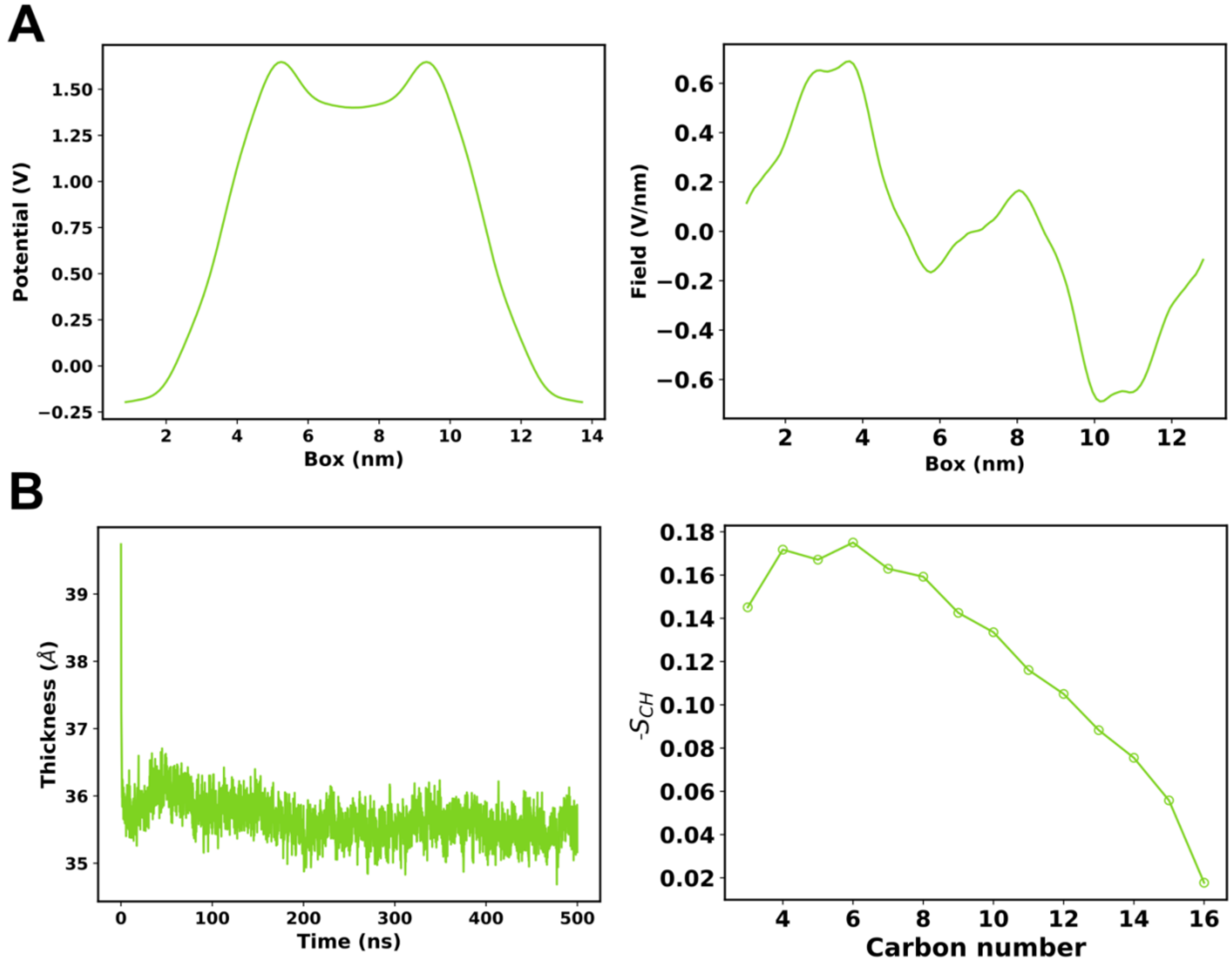


**Figure S5. A.** Potential (left) and field (right) in the system in the ΔQ 16 CompEL POPC:POPG simulation. **B.** Membrane thickness (left) throughout the 500 ns of CompEL POPC:POPG simulation, and lipid order parameters (right) of the sn-1 (palmitoyl) segment of the POPC lipid. POPC has been chosen as representative lipid to indicate the membrane ordering since it is present in all three membrane compositions.

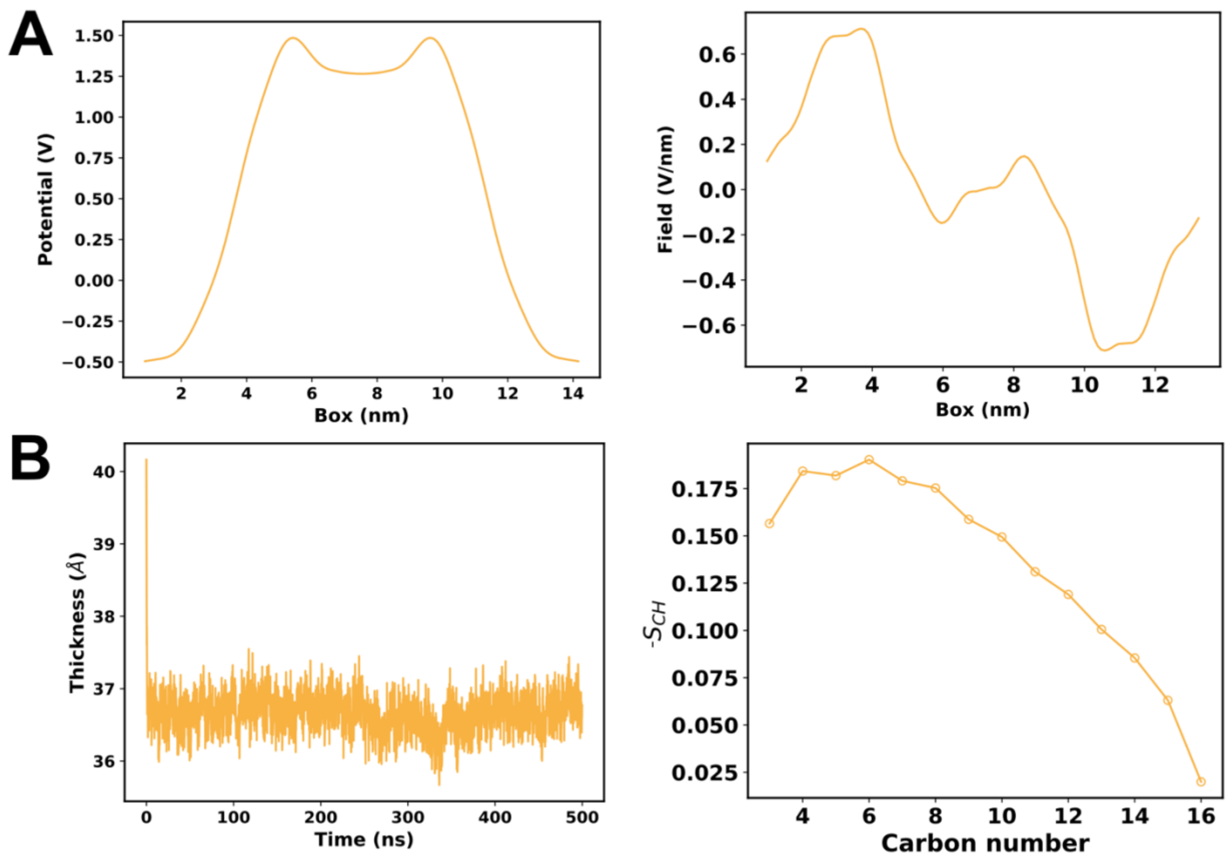


**Figure S6. A.** Potential (left) and field (right) in the system in the ∆Q 16 CompEL POPC:POPG:CHOL simulation. **B.** Membrane thickness (left) throughout the 500 ns of CompEL POPC:POPG:CHOL simulation, and lipid order parameters (right) of the sn-1 (palmitoyl) segment of the POPC lipid.

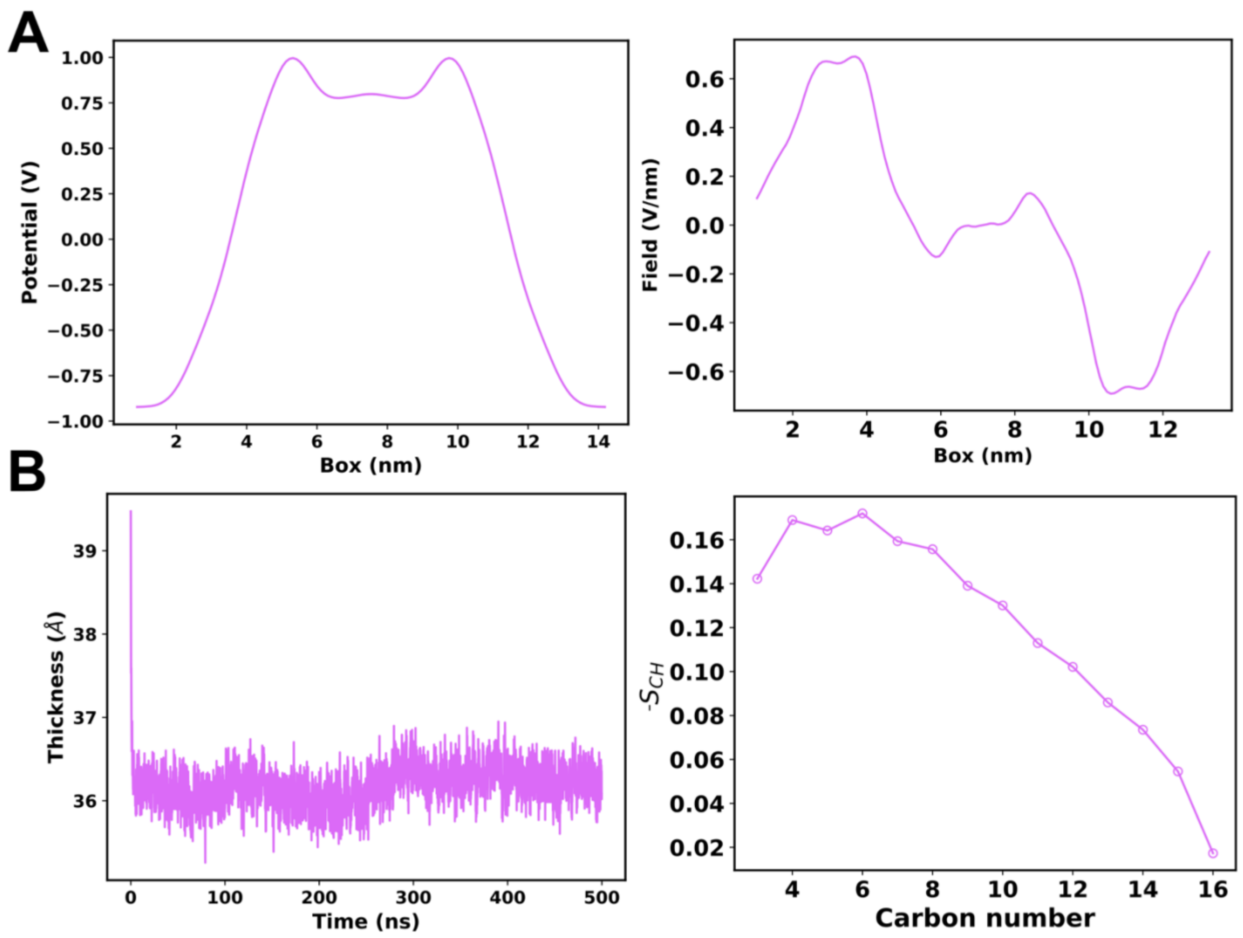


**Figure S7. A.** Potential (left) and field (right) in the system in the ΔQ 16 CompEL POPC:POPS simulation. **B.** Membrane thickness (left) throughout the 500 ns of CompEL POPC:POPS simulation, and lipid order parameters (right) of the sn-1 (palmitoyl) segment of the POPC lipid.

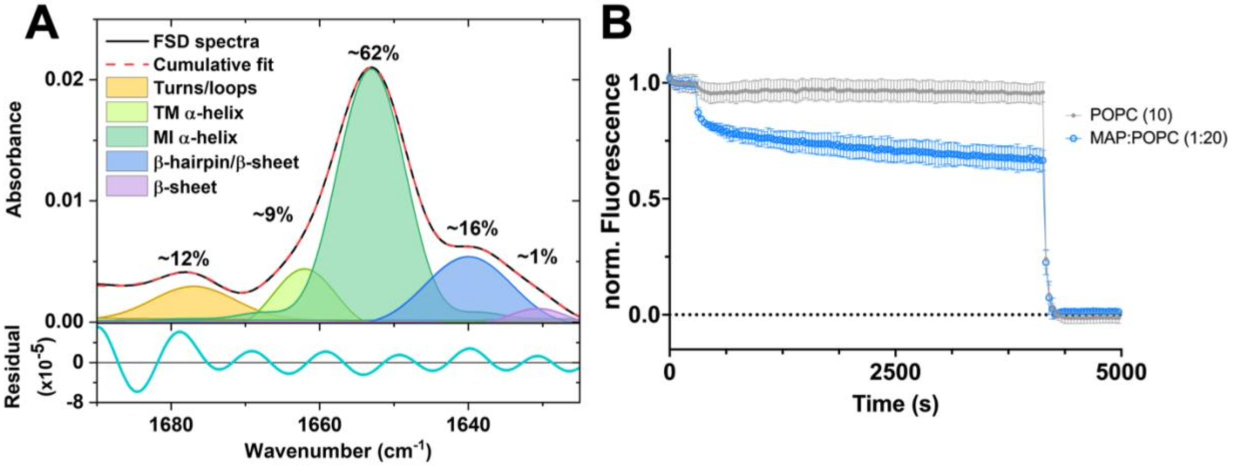


**Figure S8. A.** Average secondary structure of MAP in the FTIR experiments in POPC membranes. Mainly, the peptide adopts a membrane interface α-helical structure (~62 %, green band), followed by parallel β-sheet or β-hairpins (~16 %, blue band), turns and loops (~12 %, yellow band), transmembrane α-helix (~9 %, light green band) and β-sheet (~1 %, purple band). The fitting residual was in the order of $10^{-5}$. **B.** Liposome leakage experiments of MAP in POPC liposomes. POPC liposomes in absence (grey) and presence of MAP (blue).

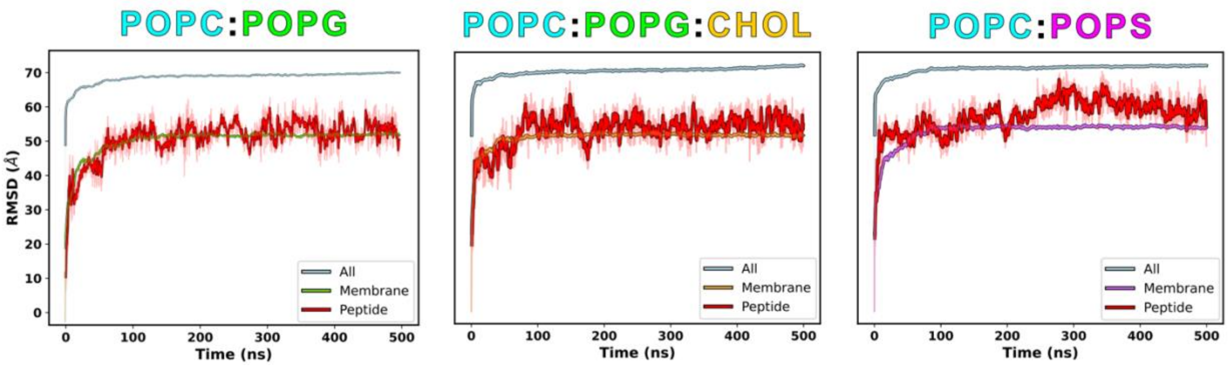


**Figure S9.** Average RMSD analysis of the CompEL ΔQ 16 simulations.

**Table S1.** Average pore radius (Å) in control simulations (without peptides).

| | POPC:POPG | POPC:POPG:CHOL | POPC:POPS |
|---|---|---|---|
| **Average** | 0.20 ± 0.02 | 0.30 ± 0.05 | 0.18 ± 0.02 |

**Table S2.** Peptide results for each CompEL ΔQ 16 simulation.

| Replica | POPC:POPG | POPC:POPG:CHOL | POPC:POPS |
|---|---|---|---|
| 1 | 8 partitioning | 1 insertion,<br>7 partitioning | 1 translocation,<br>1 insertion,<br>6 partitioning |
| 2 | 2 insertions,<br>6 partitioning | 1 insertion,<br>7 partitioning | 1 insertion,<br>7 partitioning |
| 3 | 2 insertions,<br>6 partitioning | 1 insertion,<br>7 partitioning | 1 translocation,<br>7 partitioning |

**Table S3.** Comparison of the average lipid order parameters in experimental (FTIR) and computational studies (MD CompEL simulations). Computational results of POPC simulations are taken from our previous study(Catalina-Hernandez, Aguilella-Arzo, Lopez-Martin, et al., 2025).

| Membrane | Experimental | Computational |
|---|---|---|
| POPC | -0.18 ± 0.02 | -0.12 ± 0.01 |
| POPC:POPG | -0.18 ± 0.02 | -0.12 ± 0.01 |
| POPC:POPG:CHOL | -0.19 ± 0.09 | -0.13 ± 0.01 |
| POPC:POPS | -0.19 ± 0.04 | -0.12 ± 0.01 |

**Table S4.** Comparison of averaged secondary structure estimates in experimental and computational studies. Computational results of POPC simulations are taken from our previous study(Catalina-Hernandez, Aguilella-Arzo, Lopez-Martin, et al., 2025).

| | **Experimental** | | | | | **Computational** | | |
|---|---|---|---|---|---|---|---|---|
| **Membrane** | Amide I (turns/ coiled coil structure) | Amide I (TM* α-helix) | Amide I (MI* α-helix) | Amide I (soluble peptide) | Amide I (β-sheet) | Random coil | helical | β-sheet |
| **POPC** | 11.7 | 8.75 | 62.6 | 15.7 | 1.24 | 87.9 | 3.45 | 8.69 |
| **POPC:POPG** | 11.2 | 11.6 | 60.4 | 15.2 | 1.53 | 81.1 | 4.73 | 14.2 |
| **POPC:POPG:CHOL** | 11.5 | 14.8 | 55.6 | 13.5 | 4.57 | 78.4 | 9.68 | 12.0 |
| **POPC:POPS** | 3.78 | 11.4 | 57.9 | 18.5 | 8.42 | 77.4 | 8.83 | 13.7 |
| **AVERAGE** | **9.6** | **11.6** | **59.1** | **15.7** | **4.0** | **81.2** | **6.7** | **12.1** |

*TM refers to transmembrane, and MI to membrane interface. The MI helix can be mixed with random coil contributions.
